# NRT2.1 phosphorylation prevents root high affinity nitrate uptake activity in *Arabidopsis thaliana*

**DOI:** 10.1101/583542

**Authors:** Aurore Jacquot, Valentin Chaput, Adeline Mauries, Zhi Li, Pascal Tillard, Cécile Fizames, Pauline Bonillo, Fanny Bellegarde, Edith Laugier, Véronique Santoni, Sonia Hem, Antoine Martin, Alain Gojon, Waltraud Schulze, Laurence Lejay

## Abstract

In *Arabidopsis thaliana*, *NRT2.1* codes for a main component of the root nitrate high-affinity transport system. Previous studies revealed that post-translational regulation of NRT2.1 plays an important role in the control of root nitrate uptake and that one mechanism could correspond to NRT2.1 C-terminus processing. To further investigate this hypothesis, we produced transgenic plants with truncated forms of NRT2.1. It revealed an essential sequence for NRT2.1 activity, located between the residues 494-513. Using a phospho-proteomic approach, we found that this sequence contains one phosphorylation site, at serine 501, which can inactivate NRT2.1 function when mimicking the constitutive phosphorylation of this residue in transgenic plants. This phenotype could neither be explained by changes in abundance of NRT2.1 and NAR2.1, a partner protein of NRT2.1, nor by a lack of interaction between these two proteins. Finally, the relative level of serine 501 phosphorylation was found to be modulated by nitrate in wildtype plants. Altogether, these observations allowed us to propose a model for a new and essential mechanism for the regulation of NRT2.1 activity.

## Introduction

The uptake of nitrate (NO_3_^-^) by plants from the soil solution is ensured by specific transport systems located at the plasma membrane of root cells (Krapp et al., 2014; O’Brien et al., 2016; Wang et al., 2018). In the model plant *Arabidopsis thaliana*, genes encoding root membrane NO_3_^-^ transporters have been mainly found in two separate families *NRT1* (*NPF*) and *NRT2*. In general, NRT1 proteins are low-affinity transport systems (LATS), whereas NRT2 proteins correspond to high-affinity transport system (HATS) (Miller et al., 2007; Tsay et al., 2007). To date, only NRT1.1, NRT1.2, NRT2.1, NRT2.2, NRT2.4 and NRT2.5 were shown to play a key role in the root uptake of NO_3_^-^ (Tsay et al., 1993; Huang et al., 1999; Filleur et al., 2001; Kiba et al., 2012; Lezhneva et al., 2014). However, it is clear that the HATS activity is predominantly dependent on the NRT2.1 protein. Null-mutants for *NRT2.1* have lost up to 75% of the HATS activity (Cerezo et al., 2001; Filleur et al., 2001; Li et al., 2007), and consequently cannot grow normally with NO_3_^-^ as sole N source when provided at a low concentration (e.g., <1 mM), (Orsel et al., 2004).

The regulation of *NRT2.1* has been mostly studied at the mRNA level. In particular, it has been shown that *NRT2.1* is induced upon initial NO_3_^-^ supply (Lejay et al., 1999; Girin et al., 2007) repressed by nitrogen (N) metabolites or high NO_3_^-^ provision (Lejay et al., 1999; Gansel et al., 2001; Munos et al., 2004; Krouk et al., 2006; Girin et al., 2007), and up-regulated by light and sugars (Lejay et al., 1999; Lejay et al., 2003; Lejay et al., 2008). However, previous studies suggest that in addition to transcriptional regulation, protein-protein interactions and posttranslational regulation of NRT2.1 might play an important role in modulating the activity of this NO_3_^-^ transporter. First, despite its firmly established role in root NO_3_^-^ uptake, NRT2.1 protein alone does not seem to display a NO_3_^-^ transport activity. To be functional, the Arabidopsis NRT2.1 transport system requires, like in *Chlamydomonas reinhardtii* and barley, an additional component called NAR2.1 (also called NRT3.1), a protein with a single transmembrane domain (Quesada et al., 1994; Tong et al., 2005; Okamoto et al., 2006; Orsel et al., 2006). Indeed, both gene products need to be co-expressed to yield NO_3_^-^ uptake in *Xenopus* oocytes (Orsel et al., 2006), and *nar2.1* null mutants lack the NRT2.1 protein at the plasma membrane (Wirth et al., 2007), and are strongly deficient in NO_3_^-^ HATS (Okamoto et al., 2006; Orsel et al., 2006). The precise function of NAR2.1 remains unclear, but it has been proposed that the active form of the transporter is in fact a NRT2.1/NAR2.1 hetero-oligomer (Yong et al., 2010). Second, the abundance of NRT2.1 protein in the plasma membrane shows much slower changes than those of the NO_3_^-^ HATS activity in response to light, sugars and high N supply, suggesting activation/inactivation of the NRT2.1/NAR2.1 transport system at the plasma membrane. whereas much faster changes in *NRT2.1* mRNA or NO_3_^-^ HATS activity have been demonstrated (Wirth et al., 2007; Laugier et al., 2012).

In spite of the above evidence supporting a possible posttranslational regulation of NRT2.1, the underlying mechanisms are unclear, and several hypotheses can be put forward. First, the association/dissociation dynamics of the NRT2.1/NAR2.1 hetero-dimer can correspond to such a mechanism. However, it has never been reported that this dynamics is regulated and that it actually modulates the activity of the NRT2.1/NAR2.1 transport system. Second, NRT2.1 seems to be subjected to partial proteolysis as it was shown using NRT2.1-GFP plants (Wirth et al., 2007). The possible role of this partial proteolysis is still unknown but the fact that the NRT2.1 truncated part of NRT2.1-GFP, that remains in the plasma membrane (PM), was still recognized by the anti-NRT2.1 20 antibody, which targets an epitope located 18 amino acids upstream of the C terminus of NRT2.1, suggested that the cleavage site is located in or after this epitope. This highlights a putative processing of the C-terminal part of NRT2.1 as a mechanism for controlling its activity. Third, three phosphorylation sites have been found for NRT2.1 in two phospho-proteomic approaches in response to N (Engelsberger and Schulze, 2012; Menz et al., 2016). Among them, NRT2.1 seems to be phosphorylated at S28 when plants are starved with N and rapidly dephosphorylated upon resupply of NO_3_^-^ (Engelsberger and Schulze, 2012). It suggests that, as for many membrane transport proteins, posttranslational modifications through phosphorylation are involved to control NRT2.1 activity in response to environmental cues. This conclusion fits with those of concomitant studies which demonstrated that phosphorylation of NRT2.1 ortholog in the yeast *Pichia angusta* (formerly known as *H. polymorpha*) (YNT1) is required for its delivery to the plasma membrane in response to N limitation (Navarro et al., 2008).

Another aspect, which could involve the occurrence of post-translational modifications, concerns the role of NRT2.1 in the control of root development in a way that is independent from its transport activity (Little et al., 2005; Remans et al., 2006). This gave rise to the hypothesis that NRT2.1, like NRT1.1, may also be a NO_3_^-^ sensor, or a signal transducer, but the mechanisms involved are still not known (Little et al., 2005). However, since the sensing function of NRT1.1 depends on NRT1.1 phosphorylation at T101 (Ho et al., 2009; Bouguyon et al., 2015), it supports the hypothesis that post-translational modifications of NRT2.1 could also be involved in the sensing function of NRT2.1.

To explore further and characterise the role of NRT2.1 post-translational modifications we combined the identification of NRT2.1 phosphorylation sites with the production of transgenic plants truncated in NRT2.1 C-terminus or carrying point mutations to mimic or prevent NRT2.1 phosphorylation. It allowed us to reveal the importance of the phosphorylation site S501 for NRT2.1 root NO_3_^-^ uptake activity and to propose a model for a new and essential mechanism for NRT2.1 post-translational regulation.

## Results

### NRT2.1 C-terminal part is required for root NO_3_^-^ uptake activity

Based on our previous study showing evidence for partial proteolysis of NRT2.1 at the C-terminus (Wirth et al., 2007), we investigated a putative role for NRT2.1 C-terminus on the activity of this transporter by generating transgenic plants expressing truncated forms of NRT2.1 in the *nrt2.1-2* knock-out mutant under the control of the *NRT2.1* promoter. Two different forms of NRT2.1 were produced, truncated either at the beginning (Plants ΔC_494-530_) or at the end (Plants ΔC_514-530_) of the epitope of anti-NRT2.1 20 antibody (Figure 1). For each construct, two independent transgenic lines (Δ3C_494-530_, Δ5C_494-530_) and (Δ3C_514-530_, Δ6C_514-530_) were selected based on the correct expression and regulation of the transgenes in response to NO_3_^-^ induction compared to WT plants (Figure 2C). Strikingly, the ΔC_494-530_ plants supplied with 1 mM NO_3_^-^ displayed a dramatic growth deficiency phenotype, similar to the one observed in the *nrt2.1-2* mutant (Figure 2A). On the contrary, the growth phenotype of the ΔC_514-530_ plants was similar to the WT, suggesting efficient complementation of *nrt2.1-2* mutant by the *pNRT2.1::NRT2.1ΔC_514-530_* transgene. This indicates that the C terminal part, between residues 494 and 513 and largely corresponding to the sequence of the epitope of anti-NRT2.1 20 antibody, is strictly required for correct NRT2.1 function (Figure 2A). To confirm this result, NO_3_^-^ HATS activity, determined by root ^15^NO_3_^-^ influx assays (0.2 mM external ^15^NO_3_^-^ concentration), was characterized in all lines in response to NO_3_^-^ induction, were performed (Figure 2B). In this experiment, plants were starved for N for 5 days and transferred during 4h and 7h on 1 mM NO_3_^-^. As expected, this treatment strongly stimulated both NO_3_^-^ influx (between 5 and 6-fold after 4h and 7h on 1 mM NO_3_^-^) and *NRT2.1* expression in WT plants but not in *nrt2.1-2* mutant, confirming that NO_3_^-^ HATS regulation in those conditions was mainly due to NRT2.1 activity (Figures 2B and 2C). Interestingly and in agreement with the growth phenotypes, root NO_3_^-^ influx was induced by NO_3_^-^ in ΔC_514-530_ plants almost like in the WT but not in ΔC_494-530_ plants, where it remains at a low level, similar to that in *nrt2.1-2*, while *NRT2.1* expression was normally induced in both ΔC_514-530_ and ΔC_494-530_ plants (Figures 2B and 2C). Moreover, this lack of NO_3_^-^ induction was not the only mis-regulation of root NO_3_^-^ influx observed in ΔC_494-530_ plants as responses to high N, light and sugar were also strongly altered. Indeed, in ΔC_494-530_ plants both the repression of root NO_3_^-^ influx by provision of 10 mM NH_4_NO_3_ and its induction by light or 1% sucrose were, like in *nrt2.1-2* mutant, almost abolished compared to WT and ΔC_514-530_ plants (Supplemental Figures 1A and 1B). It confirmed that the C-terminal part of NRT2.1 corresponding to 494-513 sequence is required for correct NRT2.1 activity and regulation in a large variety of experimental conditions.

**Figure 1.**
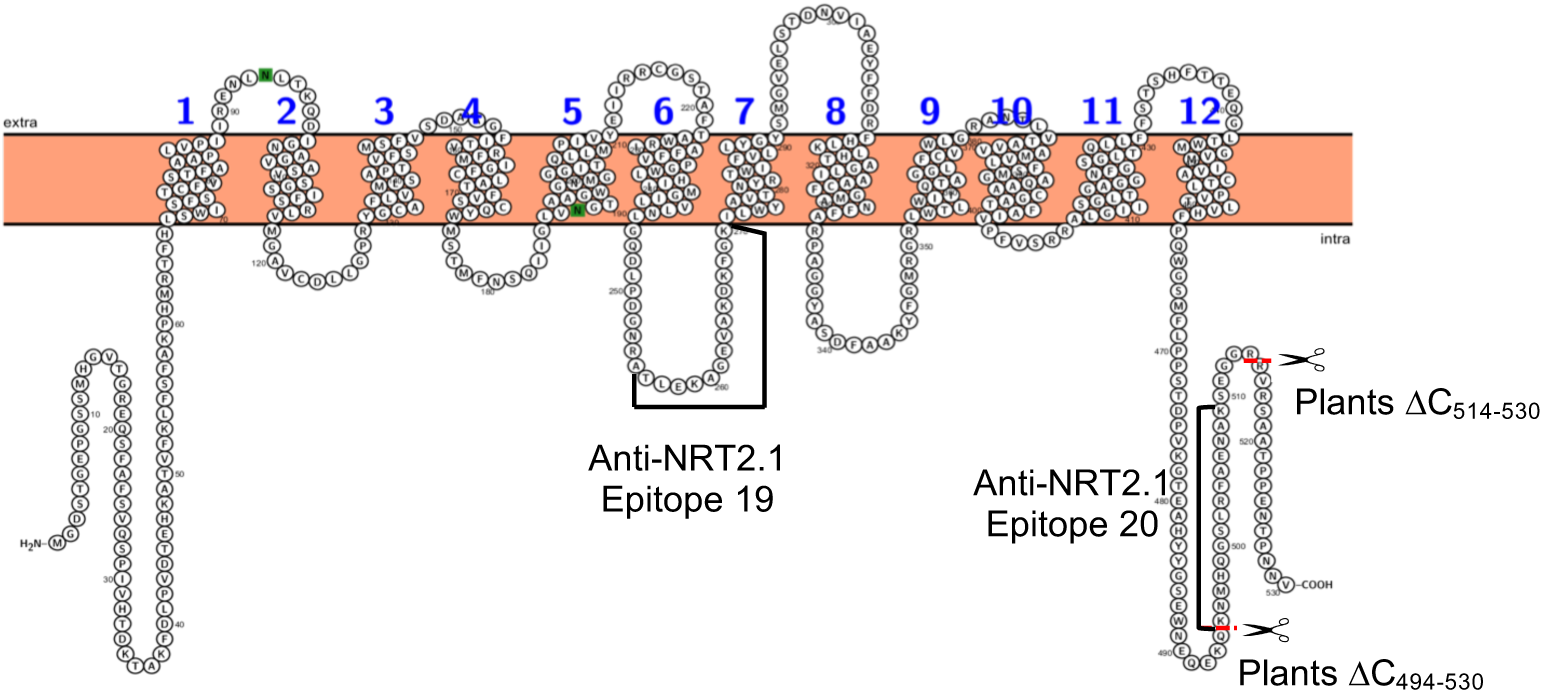
Membrane topology of NRT2.1 using Protter (Omasits et al., 2014), with the epitopes corresponding to NRT2.1 antibodies (19) and (20) and the localisation of NRT2.1 C-terminal parts removed in the plants called ΔC_494-530_ (without epitope 20) and ΔC_514-530_ (with epitope 20).

**Figure 2.**
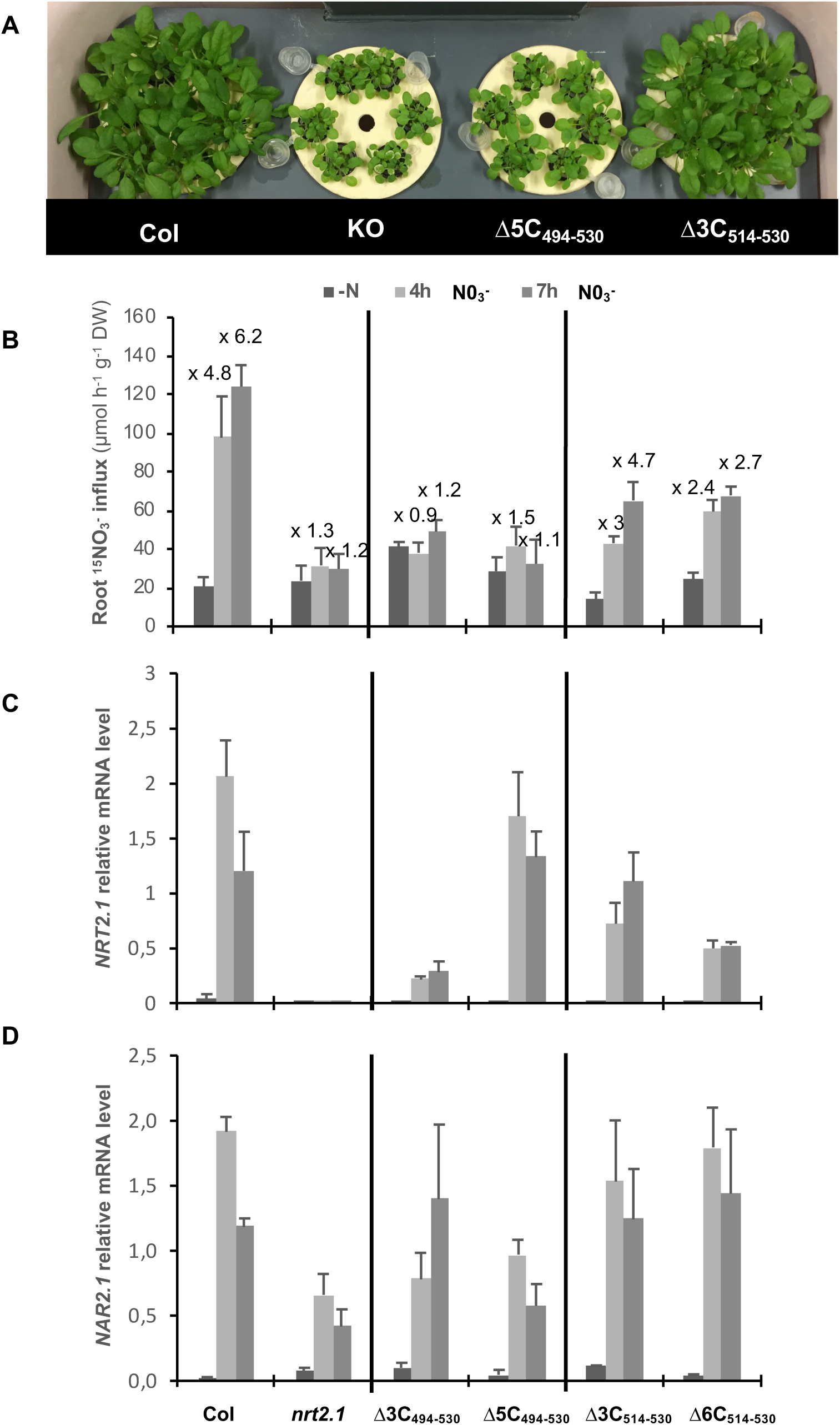
Characterization of Arabidopsis transgenic lines expressing truncated forms of NRT2.1 in the C-terminus of the protein. Wild type (Col), *nrt2.1-2* knockout mutant (*nrt2.1*) and transgenic lines expressing truncated forms of NRT2.1 without (Δ3C_494-530_ and Δ5C_494-530_) or with epitope 20 (Δ3C_514-530_ and Δ6C_514-530_). Plants were grown on 1 mM NO_3_^-^ for 5 weeks and were starved for N during 5 days. Thereafter, the plants were re-supplied with 1 mM NO_3_^-^ during 4h or 7h. **(A)** Phenotype of the plants grown on 1 mM NO_3_^-^. **(B)** Root NO_3_^-^ influx measured at the external concentration of 0.2 mM ^15^NO_3_^-^ (Values are means of 12 replicates ± SD). **(C)** and **(D)** Root *NRT2.1* and *NAR2.1* expression quantified by QPCR (Values are means of three replicates ± SD).

To further investigate the reasons of the lowered HATS activity in ΔC_494-530_ plants measurements of *NAR2.1* expression and NRT2.1 and NAR2.1 protein levels were performed in response to NO_3_^-^. Indeed, a possibility would be that in addition to a specific effect on NRT2.1 activity, NRT2.1 truncation in ΔC_494-530_ plants affects the activity and regulation of the NRT2.1/NAR2.1 complex through altered NRT2.1 and/or NAR2.1 expression and/or stability in these plants. Therefore, mRNA and PM fractions were isolated from plants starved for N for 5 days or transferred during 4h on 1 mM NO_3_^-^. The qRT-PCR and Western blot results showed that neither NAR2.1 gene expression nor its protein level were affected in ΔC_494-530_ plants compared to ΔC_514-530_ and WT plants (Figures 2D, 3A and 3B). For NRT2.1, western blots performed with Anti-NRT2.1 20 antibody confirmed that the C-terminus part containing the epitope is absent in ΔC_494-530_ plants but not in ΔC_514-530_ plants (Figure 3B). Moreover, the use of another antibody, Anti-NRT2.1 19, targeting an epitope located in an internal loop (Figure 1), revealed that NRT2.1 protein is present at similar levels in PM of ΔC_494-530_ and WT plants, which could not explained the reduction of HATS activity observed in ΔC_494-530_ plants (Figure 3A). However, it is interesting to note that in ΔC_514-530_ plants the level of NRT2.1 seems to be higher than in the WT and ΔC_494-530_ plants (Figures 3A and 3B). It suggests that the 514-530 C-terminal sequence is important for NRT2.1 stability in PM and acts in downregulating NRT2.1 protein accumulation in PM. However, this increase in NRT2.1 protein level in ΔC_514-530_ plants did not lead to an increase in root NO_3_^-^ influx compared to the WT (Figure 2B).

**Figure 3.**
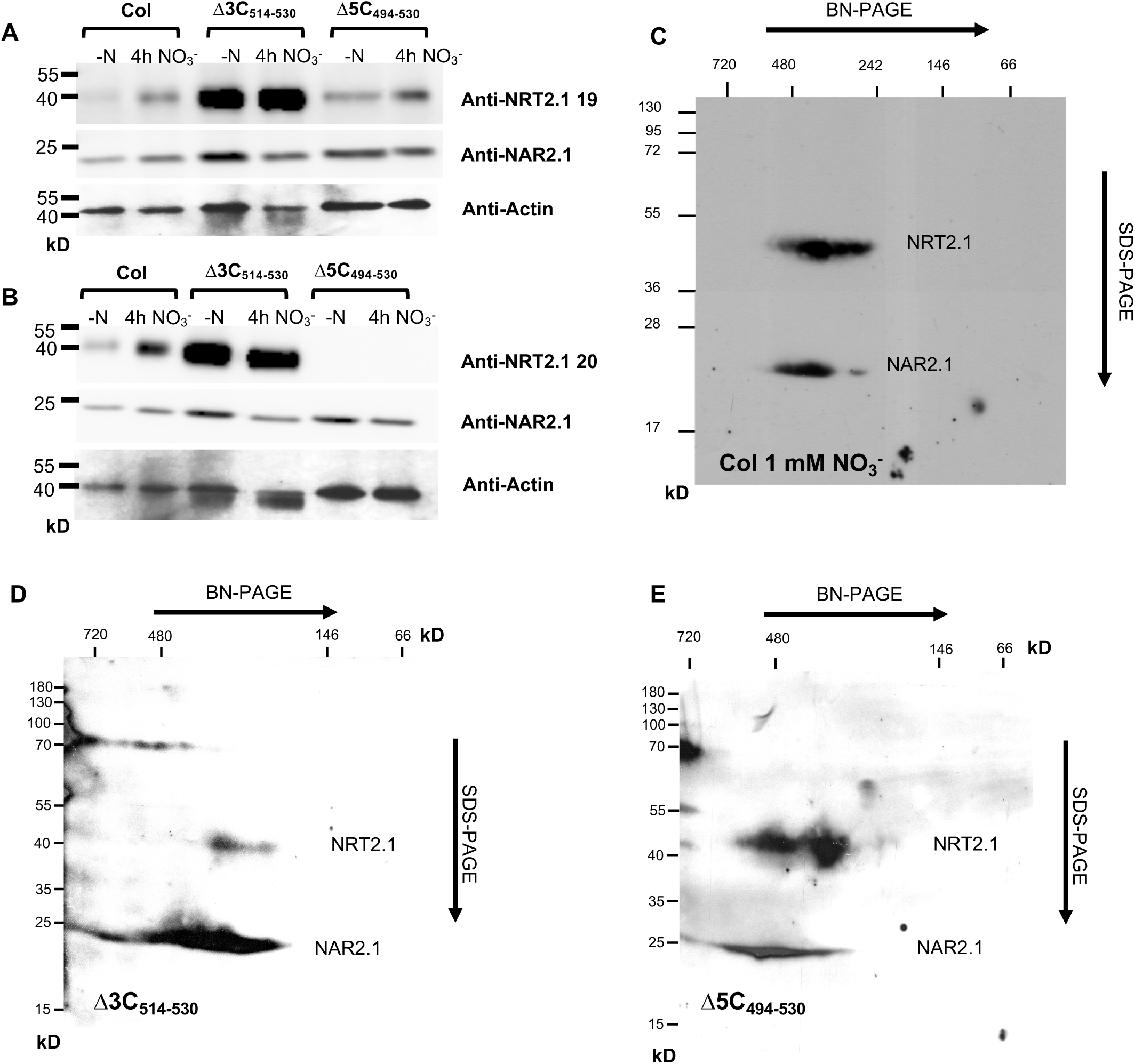
NRT2.1 and NAR2.1 protein level and protein complex in the transgenic lines expressing truncated forms of NRT2.1 in the C-terminus of the protein. **(A)** and **(B)** Immunoblot for NAR2.1 and NRT2.1 using plasma membranes extracted from roots of wild type (Col) and transgenic lines expressing truncated forms of NRT2.1 with (Δ3C_514-530_) or without epitope 20 (Δ5C_494-530_). Plants were grown on 1 mM NO_3_^-^ for 5 weeks and were starved for N during 5 days. Thereafter, the plants were re-supplied with 1 mM NO_3_^-^ during 4h. **(A)** Immunoblot for NRT2.1 using anti-NRT2.1(19) antibody. **(B)** Immunoblot for NRT2.1 using anti-NRT2.1 (20) antibody. Samples were separated on a 12% SDS-PAGE (10 µg of protein/lane). **(C)**, **(D)** and **(E)** Blue-native PAGE (BN-PAGE) for NRT2.1 and NAR2.1 complex, using microsomes extracted from roots of wild-type (Col) grown on 1 mM NO_3_^-^ **(C)** and Δ3C_514-530_ **(D)** and Δ5C_494-530_ **(E)** transgenic lines, after 4h of NO_3_- re-supply. Membranes were probed with both anti-NRT2.1 20 and anti-NAR2.1 antibodies.

Finally, to make sure that the interaction between NAR2.1 and NRT2.1 is not affected in ΔC_494-530_ plants compared to ΔC_514-530_ or WT plants, Blue native polyacrylamide gel electrophoresis (BN-PAGE) for NRT2.1 and NAR2.1 were performed using microsomes isolated from ΔC_514-530_ and ΔC_494-530_ plants induced by 1 mM NO_3_^-^ for 4h. In control WT plants grown on 1 mM NO_3_^-^ we found, as described previously (Yong et al., 2010), that NRT2.1 and NAR2.1 derived from the same protein complex (Figure 3C). However, in our case, the protein complex was approximately 400 kDa while the one described by Yong et al. (2010) was smaller, around 150 kDa. As expected in plants ΔC_514-530_, where NRT2.1 is active after an induction of 4h on 1 mM NO_3_^-^, the same ∼400 kDa protein complex with NRT2.1 and NAR2.1 was found (Figure 3D). More surprisingly, this complex was also present in plants ΔC_494-530_, where the NRT2.1/NAR2.1 complex is not active in those conditions, showing that the lack of activity is not due to a default of interaction between NRT2.1 and NAR2.1 (Figure 3E).

### Role of NRT2.1 phosphorylation in root NO_3_^-^ uptake activity

The above data indicating a key role of the C-ter 494-513 sequence of NRT2.1 in determining root NO_3_^-^ influx prompted us to make a parallel with data obtained from another approach aiming at investigating the tentative role of phosphorylation in the post-translational regulation of NRT2.1. Indeed, a mass spectrometry phosphoproteomic approach was taken to identify residues phosphorylated *in vivo*. Microsomes were isolated from WT and NRT2.1-GFP plants (Wirth et al., 2007) grown on 1 mM NO_3_^-^ and harvested in the light. Phosphorylation sites were then identified with high-accuracy mass spectrometric phosphopeptide detection using an ion trap mass spectrometer Amazon Speed ETD (Figure 4A). Using this strategy 4 phospho-peptides were identified with 2 serines phosphorylated in NRT2.1 N-terminal part (S11 and S28) and 1 serine and 1 threonine phosphorylated in NRT2.1 C-terminal part (S501 and T521) (Figures 4A and 4B). Interestingly, the phosphorylation site S501 is located in the C-ter 494-513. We therefore decided to more specifically investigate if S501 phosphorylation could affect NRT2.1/NAR2.1 activity. To do so, transgenic lines were generated by expressing point S501 mutants of NRT2.1 in the *nrt2.1-2* knock-out mutant under the control of the *NRT2.1* promoter. In the mutant versions, the S501 residue was replaced by either an alanine (A) to generate a constitutively non-phosphorylable form or with an aspartic acid (D) to mimic a constitutively phosphorylated form.

**Figure 4.**
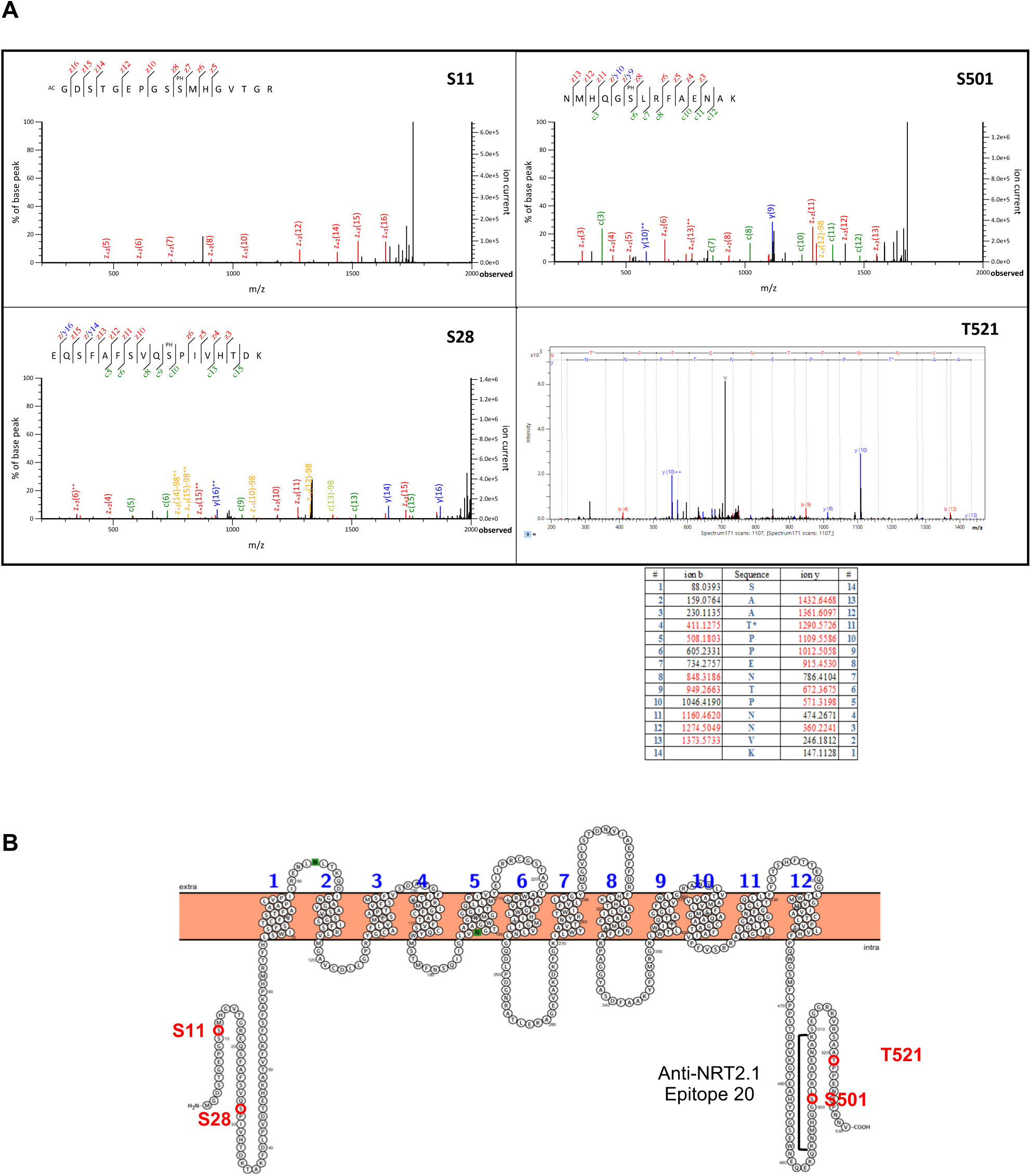
Identification of NRT2.1 phosphorylation sites. **(A)** S11: MS/MS spectrum corresponding to a serie of z ions resulting from ETD fragmentation of NRT2.1 peptide (ion score Mascot 2.6: 67; site phosphorylation probability Mascot 2.6: 96.42%). Data analysis led to the identification of the sequence GDSTGEPGSS*MHGVTGR with phosphorylation of S11. S28 and S501: MS/MS spectrum corresponding to a series of c and z ions resulting from ETD fragmentation of NRT2.1 peptides: S28: ion score Mascot 2.6: 80; site phosphorylation probability Mascot 2.6: 99.66%; S501: ion score Mascot 2.6: 54. Data analysis led to the identification of the sequence EQSFAFSVQS*PIVHTDK with phosphorylation of S28 and of the sequence NMHQGS*LRFAENAK with phosphorylation of S501. T521: MS/MS spectrum and fragmentation table corresponding to a series of b and y ions that resulted from CID fragmentation of NRT2.1-GFP peptide (ion score Mascot 2.6: 50; site phosphorylation probability Mascot 2.6: 92.69%). Data analysis led to identification of the sequence SAAT*PPENTPNNVK with phosphorylation of T521. **(B)** Membrane topology of NRT2.1 using Protter (Omasits et al., 2014), with the epitope corresponding to NRT2.1 antibody (20) and the localisation of the phosphorylation sites identified by mass spectrometry.

Like for ΔC_514-530_ and ΔC_494-530_ plants, two independent transgenic lines (S501A7, S501A9 and S501D1, S501D2) for each construct were selected, based on the correct expression and regulation of the transgenes in response to NO_3_^-^ induction compared to WT plants (Figure 5C). Again, strikingly, the same growth deficiency phenotype as for ΔC_494-530_ plants was observed for S501D plants, while the growth of S501A plants was similar to the WT when plants were grown on 1 mM NO_3_^-^ (Figure 5A). It suggested that mimicking the constitutive phosphorylation of S501 residue could resume, on its own, the phenotype observed in ΔC_494-530_ plants, when the peptide containing this residue was truncated. Accordingly, NO_3_^-^ HATS activity measurements, performed in the same conditions as described above, confirmed that the S501D substitution also prevents the induction of root NO_3_^-^ influx in response to NO_3_^-^, just like in ΔC_494-530_ plants, while the S501A substitution leads to the same level of root NO_3_^-^ influx as in WT plants (Figure 5B). Since in all the genotypes the level of *NAR2.1* expression was the same as in the WT and that *NRT2.1* expression was normally induced by NO_3_^-^ in both S501A and S501D plants compared to WT, default of HATS activity in S501D plants could not be explained by a default in the expression of *NRT2.1* or *NAR2.1* (Figures 5C and 5D). Furthermore, like in ΔC_494-530_ plants, the lack of root NO_3_^-^ influx due to the S501D substitution was not specific of the regulation by NO_3_^-^ and was also observed in response to high N, light and sugar (Supplemental Figures 1C and 1D). These results indicate that phosphorylation of the S501 residue has a general impact on NRT2.1 activity independently of the environmental conditions.

**Figure 5.**
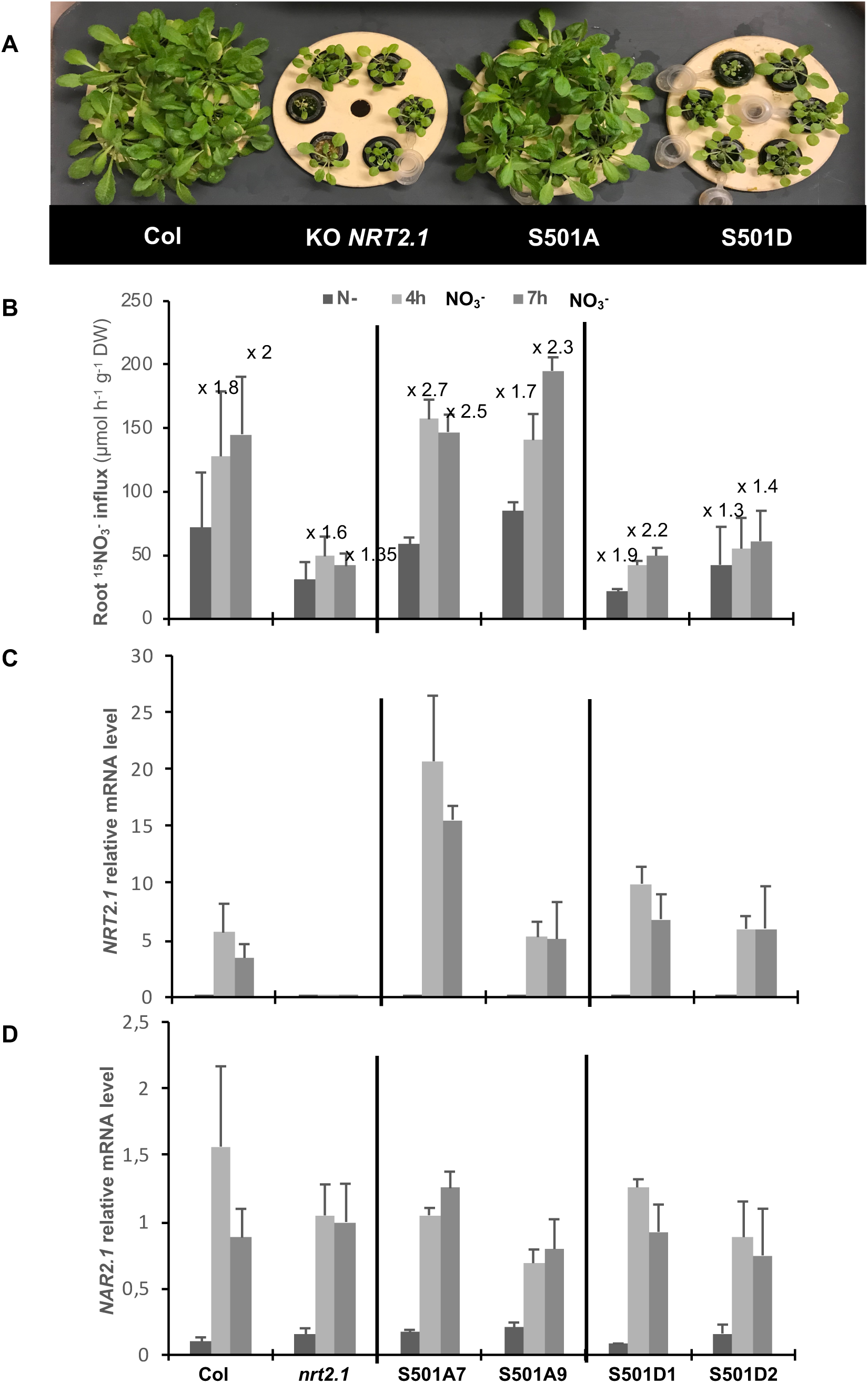
Characterization of Arabidopsis transgenic lines expressing mutated forms of NRT2.1 on the phosphorylation site S501. Wild type (Col), *nrt2.1-2* knockout mutant (*nrt2.1*) and transgenic lines expressing mutated forms of NRT2.1, which either cannot be phosphorylated (S501A7 and S501A9) or mimic a constitutive phosphorylation of S501 (S501D1 and S501D2). Plants were grown on 1 mM NO_3_^-^for 5 weeks and were starved for N during 5 days. Thereafter, the plants were re-supplied with 1 mM NO_3_^-^ during 4h and 7h. **(A)** Phenotype of the plants grown on 1 mM NO_3_^-^. **(B)** Root NO_3_^-^ influx measured at the external concentration of 0.2 mM ^15^NO_3_^-^ (Values are means of 12 replicates ± SD). **(C)** and **(D)** Root *NRT2.1* and *NAR2.1* expression quantified by QPCR (Values are means of three replicates ± SD).

Western blots and BN PAGE were then performed, using respectively PM and microsomes isolated from plants starved for N during 5 days and induced by 1 mM NO_3_^-^ during 4h. The results showed that both NAR2.1 and NRT2.1 proteins are present in all the genotypes. However, the amount of NRT2.1 protein seems to be lower in S501D plants compared to S501A and WT plants (Figures 6A and 6B). It is interesting to note that this result was obtained with both antibodies Anti-NRT2.1 (19) and (20), suggesting that this lower level of NRT2.1 in S501D plants was not due to any processing of NRT2.1 C-terminal part because of the mutation. Nevertheless, this cannot fully explain, the total lack of NRT2.1 activity in S501D plants. Indeed, NO_3_^-^ still markedly increases the level of NRT2.1 protein in those plants and should have led to a higher increase in root NO_3_^-^ influx than in the *nrt2.1-2* mutant, which was not observed. Furthermore, BN PAGE indicates that the S501D substitution has no impact on the NRT2.1/NAR2.1 protein complex, which is still found at ∼ 400 kDa in both S501A and S501D plants (Figures 6C and 6D). In this case, the normal interaction between NRT2.1 and NAR2.1 was also confirmed by rBiFC experiments performed on tobacco leaves (Figure 6E and Supplemental Figure 2). The quantification of the fluorescence signal shows that when the leaves are transformed with the WT forms of NAR2.1 and NRT2.1, YFP fluorescence is about 40% of the control RFP fluorescence, while in the leaves transformed with both NAR2.1 and NRT2.1 S501A or S501D mutated forms, YFP fluorescence is even slightly higher than with the WT form, with an YFP fluorescence of around 50% of RFP fluorescence (Figure 6E). It clearly confirms that the lack of activity of NRT2.1 S501D mutated form can be explained neither solely by a decrease in NRT2.1 or NAR2.1 protein level nor by a default of interaction between NRT2.1 and NAR2.1 in the same protein complex as the one observed in WT and in S501A plants.

**Figure 6.**
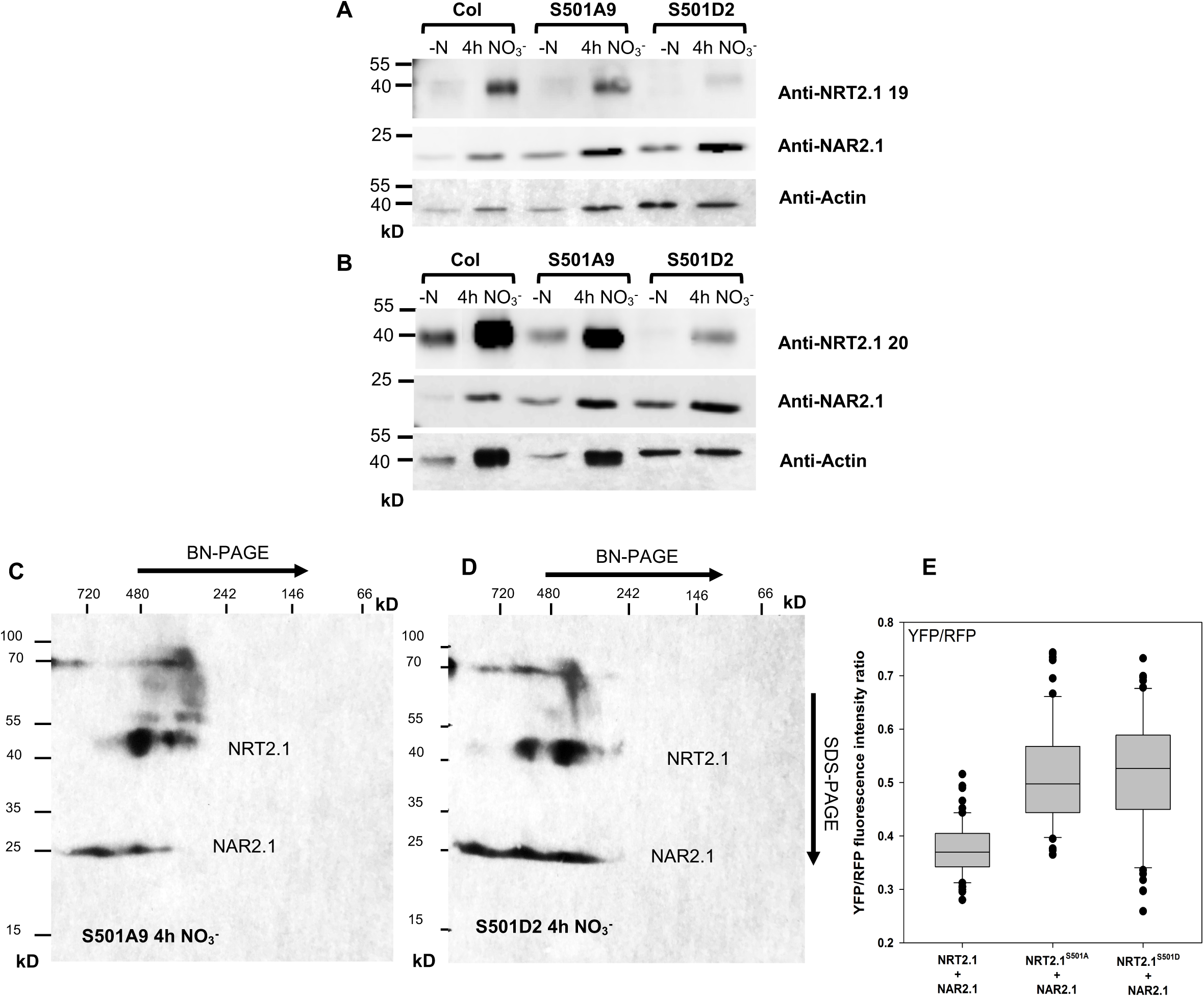
NRT2.1 and NAR2.1 protein level and protein complex in the transgenic lines expressing mutated forms of NRT2.1 for S501 phosphorylation site. **(A)** and **(B)** Immunoblot for NAR2.1 and NRT2.1 using plasma membranes extracted from roots of wild type (Col) and transgenic lines expressing mutated forms of NRT2.1, which either cannot be phosphorylated (S501A9) or mimic a constitutive phosphorylation on S501 (S501D2). Plants were grown on 1 mM NO_3_^-^ for 5 weeks and were starved for N during 5 days. Thereafter, the plants were re-supplied with 1 mM NO_3_^-^ during 4h. **(A)** Immunoblot for NRT2.1 using anti-NRT2.1(19) antibody. **(B)** Immunoblot for NRT2.1 using anti-NRT2.1 (20) antibody. Samples were separated on a 12% SDS-PAGE (10 µg of protein/lane). **(C)** and **(D)** Blue-native PAGE (BN-PAGE) for NRT2.1 and NAR2.1 complex, using microsomes extracted from roots of S501A9 **(C)** and S501D2 **(D)** transgenic lines, after 4h of NO_3_- re-supply. Membranes were probed with both anti-NRT2.1 (20) and anti-NAR2.1 antibodies. **(E)** rBiFC fluorescence signals from five independent experiments using tobacco plants transformed with the pBiFCt-2in1-CC vector. Each box plot represents the mean of fluorescence intensity ratios from more than 12 images taken at random over the leaf surface. rBiFC signals were calculated as the mean fluorescence intensity ratio of YFP to RFP determined from each image set.

### Regulation of S501 phosphorylation in WT plants

The phenotype of the S501D plants suggests that the phosphorylation of the S501 residue may correspond to a major regulatory mechanism able to inactivate the NRT2.1/NAR2.1 transport system. This raised the question of whether this phosphorylation actually occurs in vivo, especially when the HATS activity is modulated in response to environmental conditions. To follow the level of S501 phosphorylation in WT plants, two specific polyclonal antibodies, called Anti-S501P and Non-specific NRT2.1 antibody, were raised in rabbit against a NRT2.1 S501 phosphopeptide and its non-phosphorylated counterpart, respectively. The affinity-purified anti-NRT2.1 antibodies were tested with dot-blots using the synthetic peptides produced to purify the antibodies. It confirmed that Anti-S501P antibody could only recognize S501 phosphorylated peptide while NRT2.1 non-specific antibody recognized both S501 phosphorylated and non-phosphorylated peptides (Supplemental Figure 3).

To follow the level of S501 phosphorylation, WT plants were first, as described above, starved for N for 5 days and then transferred on 1 mM NO_3_^-^ during 4h. However, in those conditions, the amount of NRT2.1 after 5 days on 0N was too low to allow detection of the phosphorylated form of NRT2.1 with anti-S501P antibody using either microsomes or PM (data not shown). In order to follow the impact of NO_3_^-^ induction on S501 phosphorylation, microsomes were thus isolated from plants induced during 1h or 4h with 1 mM NO_3_^-^ after 5 days of N starvation. In those conditions, root NO_3_^-^ influx was still induced about 2-fold between 1 h and 4 h in WT plants compared to *nrt2.1-2* mutant, and the level of NRT2.1 was already high enough at 1 h to allow the phosphorylated form to be detected with S501 specific antibody (Supplemental Figures 4 and 7). Quantitative analysis by ELISA, using both anti-S501P and anti-NRT2.1-20 antibodies, revealed that, after 4h of NO_3_^-^ re-supply, the overall level of NRT2.1 protein increased, while the level of NRT2.1 S501 phosphorylated form did not change compared to plants re-supplied with NO_3_^-^ for 1h (Figure 7B). This result is consistent with a role of S501 phosphorylation in NRT2.1 inactivation as it indicates that the level of S501 phosphorylation in WT plants decreases in conditions where NRT2.1 becomes more active. But surprisingly, when western blots were performed on the same experiments, the abundance of the band corresponding to NRT2.1 at ∼ 45 kDa increased both after detection with anti-S501P and anti-NRT2.1-20 antibodies, which did not correspond to the quantitative results obtained with ELISA (Figure 7A). However, as described in Wirth et al. (2007) when using microsomes, two other bands, at ∼ 100 kDa and ∼ 130 kDa, corresponding presumably to NRT2.1 multimeric forms, were detected with both antibodies. The use of *nrt2.1-2* knock-out mutant confirmed that all the bands were specific for NRT2.1 protein with both anti-S501P and anti-NRT2.1-20 antibodies. Interestingly, NRT2.1 S501 phosphorylated form was more abundant in the high molecular mass bands after 1h of NO_3_^-^ re-supply compared to 4h, while in the same samples the level of total NRT2.1 protein in the high molecular mass bands was not affected or even slightly increased after 4h of NO_3_^-^ re-supply compared to 1h (Figure 7A). These results suggest that the apparent increase of the band at ∼ 45 kDa, for the S501 phosphorylated fraction of NRT2.1, was not due to an increase in the amount of NRT2.1 phosphorylated form in response to NO_3_^-^ but rather to a removal of the phosphorylated form of NRT2.1 from the multimeric forms of NRT2.1 in response to NO_3_^-^. In the same time, it means that the increase in total NRT2.1 protein, detected with anti-NRT2.1 20 after 4h of NO_3_^-^ re-supply, corresponds mainly to an increase in NRT2.1 non phosphorylated on S50l, which becomes the main form in the high molecular mass band after 4h of NO_3_^-^ re-supply compared to 1h (Figures 7A and 7B).

**Figure 7.**
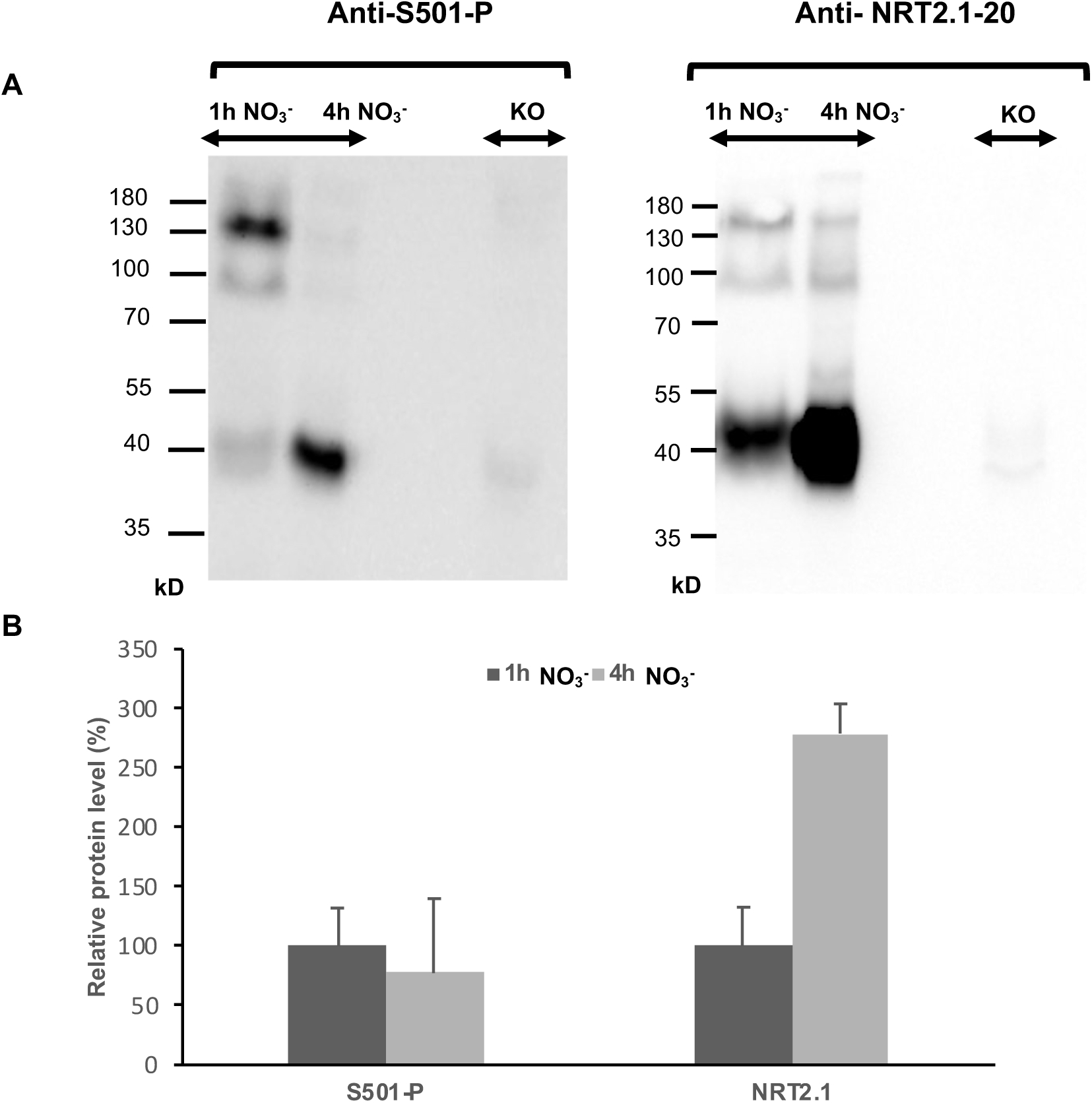
Regulation of NRT2.1 S501 phosphorylation in response to NO_3_^-^. Wild type plants were grown on 1 mM NO_3_^-^ for 5 weeks and were starved for N during 5 days. Thereafter, the plants were re-supplied with 1 mM NO_3_^-^ during 1h or 4h. Knock-out plants for *NRT2.1* (KO) were grown on 1 mM NO_3_^-^ **(A)** Immunoblot for NRT2.1 S501 phosphorylation site (Anti-S501-P) and NRT2.1 (Anti-NRT2.1-20) using microsomes extracted from roots. Samples were separated on a 12% SDS-PAGE (20 µg of protein/lane). **(B)** ELISA from three independent experiments for NRT2.1 phosphorylation site (S501-P) and NRT2.1 using microsomes extracted from roots.

### Impact of S501 mutation on root development

Previous studies showed that, independently of its role in root NO_3_^-^ uptake, NRT2.1 is involved in lateral root (LR) development (Little et al., 2005; Remans et al., 2006). It leads to the hypothesis that, like NRT1.1, NRT2.1 could be a NO_3_^-^ sensor (Krouk et al., 2010). However, the mechanism involved remains completely unknown and a key evidence to support a role for NRT2.1 in NO_3_^-^ sensing would be to uncouple its activity as a root NO_3_^-^ transporter from its role in LR development. Since the S501D substitution was able to inactivate NRT2.1 as a NO_3_^-^ transporter, it prompted us to investigate its impact on lateral root development. To determine whether lateral root development is affected by S501 phosphorylation, the total numbers of initiated LR primordia and visible LRs were scored in the newly formed portion of the primary root in WT, *atnrt2.1-2*, S501A and S501D plants after transfer from 1 mM NO_3_^-^ to 0N, 0.3 mM NO_3_^-^ or 5 mM NO_3_^-^ (Figure 8).

**Figure 8.**
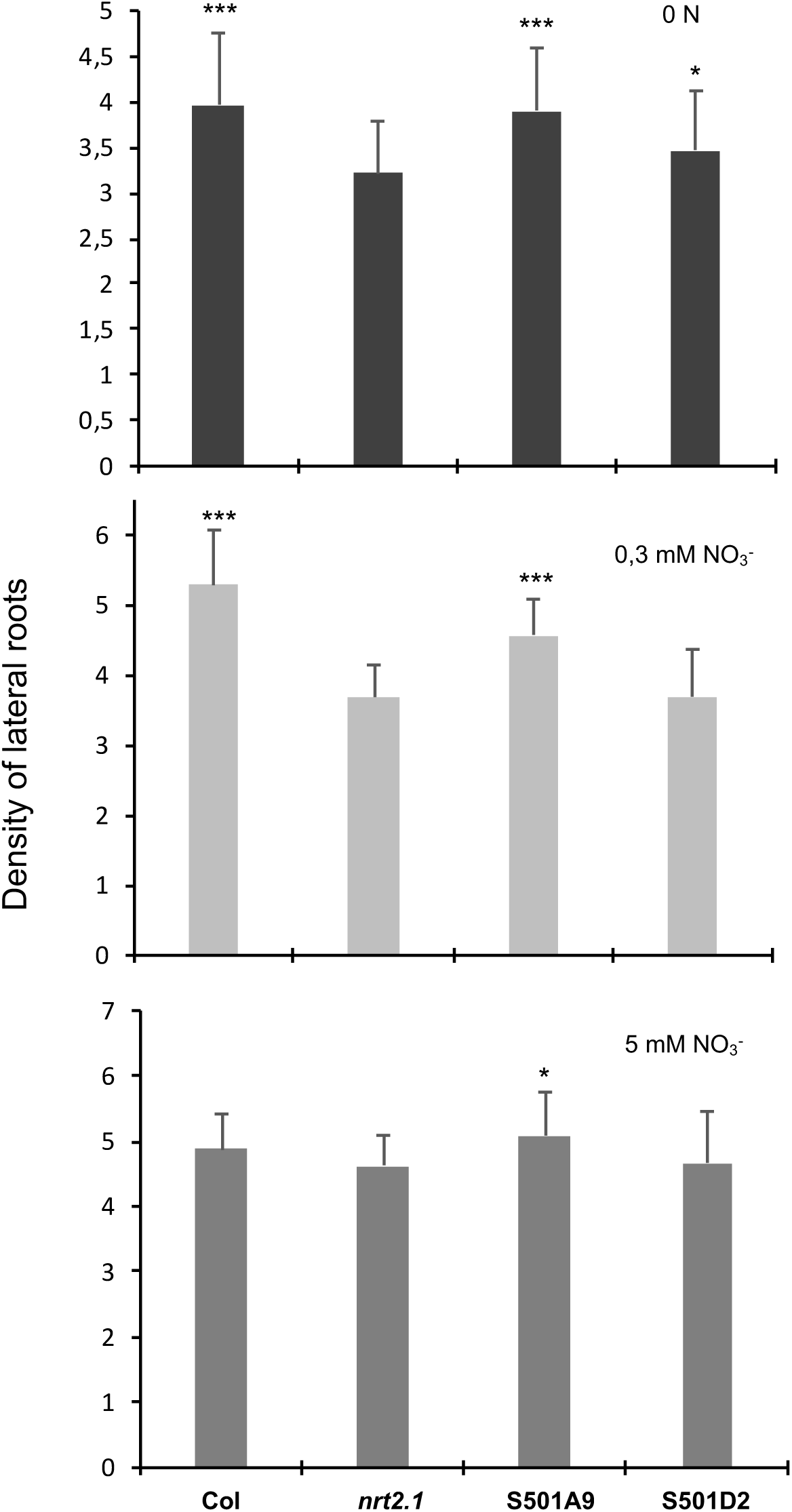
Impact of S501A and S501D mutations on lateral root density in response to NO_3_^-^ Density of initiated primordia corresponding to the total number of primordia plus lateral roots normalized by the length of the primary root of wild type (Col), *nrt2.1-2* knockout mutant (*nrt2.1*) and S501A9 or S501D2 transgenic lines. Plants were transferred at day 5 from 1 mM NO_3_^-^ to either nitrogen-free medium (0 N) or 0,3 and 5 mM NO_3_^-^. Total number of visible LRs and LR primordia were determined between day 10 and 12. Average values are means of at least 20 replicates ± SD. *P < 0.05, **P < 0.01 and ***P < 0,001 calculated by unpaired Sudents t-test compared to *nrt2.1*.

As described in Remans et al. (2006), KO mutation of *NRT2.1* resulted in a reduced LR initiation in the portion of the primary root developing after transfer to N-free medium or to the 0.3 mM NO_3_^-^ medium as compared to WT (Figure 8). This difference between the two genotypes was not observed after transfer to the 5 mM NO_3_^-^ medium, confirming that mutation of *NRT2.1* has no significant consequence on LR growth under non-limiting N supply (Orsel et al., 2004; Little et al., 2005; Remans et al., 2006). S501D plants had the same phenotype than *nrt2.1-2* knock-out plants, with a similar decrease in the density of initiated LR primordia after transfer to 0N and 0.3 mM NO_3_^-^ and no significant difference on 5 mM NO_3_^-^ compared to the WT (Figure 8). Conversely, the density of initiated LR primordia in S501A plants was similar to the WT in all the conditions. These data confirm that NRT2.1 acts as an activator of LR primordia initiation, even when its substrate is absent. They also clearly indicate that the S501D substitution is able to inactivate both NRT2.1 root NO_3_^-^ uptake activity and its role in LR development under limiting N supply, whereas the S501A substitution did not affect both NRT2.1 functions.

## Discussion

### Role of NRT2.1 phosphorylation in root NO_3_^-^ uptake activity

In previous studies we provided evidence for post-translational regulations of the root NRT2.1/NAR2.1 NO_3_^-^ transporter (Wirth et al., 2007; Laugier et al., 2012). One of the hypotheses was that NRT2.1 C-terminus was cleaved and that this mechanism could play a role in the regulation of NRT2.1 activity (Wirth et al., 2007). The results obtained with truncated forms of NRT2.1 show that ΔC_494-530_ plants have the same phenotype than *nrt2.1-2* knock-out mutant for growth and root NO_3_^-^ influx, while ΔC_514-530_ plants are like the wild-type (Figure 2). It thus seems at first sight to support the hypothesis that partial proteolysis of NRT2.1 C-terminus sequence may play a role in NRT2.1 activity. However, the characterization of NRT2.1 phosphorylation sites provides a different explanation for the phenotype observed in ΔC_494-530_ plants. We found 4 phosphorylated residues in NRT2.1: S11, S28, S501 and T521 (Figure 4). These results are consistent with previous studies showing that S11, S28 and T521 are phosphorylated under NO_3_^-^ conditions (Engelsberger and Schulze, 2012; Menz et al., 2016), but to our knowledge, this is the first time that S501 was found to be phosphorylated. Interestingly, S501 is also the only phosphorylation site identified in the C-terminus part of NRT2.1, which we found to be strictly required for NO_3_^-^ HATS activity. The phenotype of the transgenic plants we produced with NRT2.1 S501 point mutants strikingly revealed that mimicking the constitutive phosphorylation in S501D plants is able to resume the phenotype observed in ΔC_494-530_ plants (Figure 5). It indicates that the impact of NRT2.1 ΔC_494-530_ deletion on root NO_3_^-^ uptake activity is not directly due to the lack of the deleted sequence but rather to the fact that the deletion in ΔC_494-530_ plants removes S501 phosphorylation site, while it remains intact in ΔC_514-530_ plants. Indeed, the phenotype of S501D plants cannot be explained by any NRT2.1 C-terminus processing due to the substitution since NRT2.1 can still be detected in Western blots by the antibody NRT2.1 20, which targets an epitope localized specifically in the truncated part responsible for NRT2.1 inactivation in ΔC_494-530_ plants (Figures 6A and 6B).

Furthermore, the fact that S501A has the same phenotype as wild type plants indicates that S501 phosphorylation is likely involved in NRT2.1 inactivation, while S501 dephosphorylation enables NRT2.1 activation. It is also interesting to note that the loss of NRT2.1 activity in S501D plants is not only observed in response to NO_3_^-^ induction. Indeed, the lack of root NO_3_^-^ uptake activity in ΔC_494-530_ and S501D plants is also observed in response to induction by light and sugar and repression by 10 mM NH_4_NO_3_ (Supplemental Figure 1). It supports the hypothesis that the absence of phosphorylation of NRT2.1 S501 residue is required for NRT2.1 activity and that it cannot be overcome by any other post-translational modifications potentially triggered by other environmental factors.

### Characterization of the mechanisms involved in NRT2.1 inactivation

Since phosphorylation can induce changes in protein activity (Liu and Tsay, 2003), provide docking sites for protein-protein interaction (Pawson and Scott, 1997) or induce changes in subcellular localization (Navarro et al., 2008), we tested the impact of S501 mutations on those different aspects in order to further characterize the mechanisms involved in NRT2.1 inactivation.

The fact that NRT2.1 is still detected in ΔC_494-530_ or S501D plants, when western blots are performed with purified plasma membrane fractions, ruled out the hypothesis that the complete lack of NRT2.1 activity in those plants is due to changes in its subcellular location compared to WT plants (Figures 3A, 3B, 6A and 6B).

Protein-protein interaction is a known mechanism for NRT2.1 since it requires a second protein, NAR2.1 (Okamoto et al., 2006; Orsel et al., 2006), to generate a heterooligomer that may be the active form of the transporter (Yong et al., 2010). However, to date, the mechanism remains unknown and S501 phosphorylation could prevent NRT2.1 transport activity by disrupting NRT2.1 and NAR2.1 interaction. Therefore, we performed BN-PAGE and rBiFC experiments to test the interaction between NAR2.1 and NRT2.1 S501A and S501D mutated forms compared to WT form (Figures 6C, 6D and 6E). The data show that, even when NRT2.1 is inactive in S501D plants, the mutation does not change the size of the protein complex at ∼ 480 kDa nor the interaction with NAR2.1 inside this complex. These results indicate that S501 phosphorylation does not prevent NRT2.1 interaction with NAR2.1 to explain the loss of NRT2.1 activity in S501D plants compared to S501A and WT plants. This conclusion is supported by the BN-PAGE experiments performed with the truncated forms of NRT2.1 C-terminus part (Figures 3C, 3D and 3E). Indeed, the size of the complex at ∼ 480 kDa and the presence of NAR2.1 in this complex is the same in ΔC_494-530_ plants, where NRT2.1 is not active, compared to ΔC_514-530_ and WT plants, where NRT2.1 is active. It confirms that neither S501 phosphorylation states nor NRT2.1 C-terminus region, from at least the amino acids 493 to 530, can provide a docking site for NAR2.1 interaction. The conclusion that NRT2.1 C-terminus is not involved in the interaction with NAR2.1 is supported by the recent yeast two-hybrid (Y2H) experiments performed by Kotur et al. (2017). In this work, the authors identified a leucine residue located in the first putative trans-membrane region at position 85 of NRT2.1, which when mutated to glutamine resulted in disruption of the interaction between NAR2.1 and NRT2.1. Finally, it has been shown previously that in the *nar2.1* mutants, NRT2.1 protein is absent although mRNA encoding NRT2.1 is present (Wirth et al., 2007; Yong et al., 2010). It leads to the hypothesis that, unless NAR2.1 is available to generate the NAR2.1/NRT2.1 complex, the NRT2.1 protein is degraded. Thus, if NRT2.1/NAR2.1 interaction was affected in ΔC_494-530_ and S501D plants, we should have observed an absence of NRT2.1 protein in those plants.

If disruption of protein-protein interaction between NRT2.1 and NAR2.1 cannot explain the phenotype of ΔC_494-530_ and S501D plants, it cannot be ruled out that the removal of NRT2.1 C-terminus containing S501, in ΔC_494-530_ plants, or S501D substitution are able to prevent interaction with other unknown partner proteins involved in NRT2.1 activity. Indeed, the protein complex we detect is very large (∼ 480kDa) and could contain several subunits of NRT2.1 and NAR2.1 associated with other proteins. In Yong et al. (2010) however, the authors found a protein complex of only ∼ 150kDa, that they explained by the interaction of two subunits each of NRT2.1 and NAR2.1. The reason for this discrepancy in the size of the protein complex between our results and theirs are not clear. Perhaps the buffers and/or conditions used for isolation of microsomes/plasma membranes and/or BN-PAGE slightly differ between our two laboratories leading to partial denaturation of a bigger NRT2.1/NAR2.1 protein complex in Yong et al. (2010) conditions. More denaturing conditions could also explain why Yong et al. (2010) do not detect any anti-NRT2.1 reactive polypeptides at approximately 75 and 120 kDA in regular Western-blots as it was found by Wirth et al. (2007). This would support the hypothesis of a core complex constituted of two or more subunits each of NRT2.1 and NAR2.1 associated with one or several proteins, which could be more sensitive to partial denaturation depending on the strength of their interactions with NRT2.1 and/or NAR2.1.

### S501 phosphorylation in wild-type plants

The results obtained with the transgenic plants expressing NRT2.1 S501 point mutants raised the question of the regulation of S501 phosphorylation in wild-type plants. If mimicking constitutive phosphorylation of S501 leads to inactivation of NRT2.1, one would expect that in wild-type plants, NRT2.1 S501 residue is phosphorylated in conditions where NRT2.1 is not active and not phosphorylated when plants are transferred in conditions where NRT2.1 is activated. In Arabidopsis, inactivation of a transporter through phosphorylation of the C-terminus part of the protein has already been demonstrated for the ammonium transporter AMT1.1 (Loque et al., 2007). Indeed, AMT1.1 in plants, like MEP2 in yeast, works as a trimer, which activity is controlled by the spatial positioning of the C-terminus. When ammonium is added in the growth medium, it triggers rapid phosphorylation of a conserved threonine residue (T460) in the C-terminus of AMT1.1 in a time- and concentration-dependent manner (Loque et al., 2007; Lanquar et al., 2009). This phosphorylation of T460 in response to an increase in external ammonium correlates with a reduction of ammonium uptake activity in roots. These results lead to a model in which T460 phosphorylation induces a conformational change and that a single phosphorylation event in the C-terminus of one monomer is sufficient for cooperative closure of the trimer (Lanquar et al., 2009; Lanquar and Frommer, 2010).

For NRT2.1, the ELISA data show that NRT2.1 S501 phosphorylated form is relatively less abundant compared to NRT2.1 unphosphorylated form after 4h of NO_3_^-^ re-supply (Figure 7B). It correlates with an induction of root NO_3_^-^ uptake activity in the same experiment (Supplemental Figure 4) and thus support the hypothesis that when NO_3_^-^ is added in the medium, it triggers S501 de-phosphorylation, which activates NRT2.1. However, ELISA experiments also show that the level of NRT2.1 protein increases about 2.5 fold after 4h of NO_3_^-^ induction compared to 1h, while in the same time the amount of NRT2.1 S501 phosphorylated form is not affected by NO_3_^-^ re-supply. It suggests that in fact, NO_3_^-^ does not activate NRT2.1 de-phosphorylation but rather induces synthesis of new unphosphorylated NRT2.1 protein. Furthermore, compared to studies on ammonium, the kinetics of root NO_3_^-^ uptake induction after NO_3_^-^ re-supply is rather slow. Indeed, when N-deficient Arabidopsis plants are resupplied with ammonium, ammonium influx into roots is repressed within minutes (Rawat et al., 1999), while it takes several hours for root NO_3_^-^ uptake to be induced after NO_3_^-^ resupply (Supplemental Figure 4) (Cerezo et al., 2001). These data also support the idea that NRT2.1 activation after NO_3_^-^ resupply is not due to rapid S501 dephosphorylation.

The Western blots we performed on the same samples used for ELISA revealed an even more complex picture. Indeed, both the level of NRT2.1 and NRT2.1 S501 phosphorylated form, detected at ∼ 45 kDa, were induced after 4h of NO_3_^-^ resupply compared to 1h (Figure 7A). This result was very surprising since it did not reconcile with the ELISA experiments showing only an increase of NRT2.1 protein level after 4h of NO_3_^-^ resupply and not of the phosphorylated form (Figure 7B). However, when increasing the exposure time of the membranes, we were able to detect with both S501-P and NRT2.1-20 antibodies two other bands with a higher molecular weight at ∼ 100 kDa and ∼ 150 kDa. These bands have already been described by Wirth et al. (2007) as specific of NRT2.1 and as revealing higher molecular mass complexes incorporating NRT2.1. The fact that NRT2.1 is part of a high molecular weight complex together with NAR2.1 has been confirmed in this study and by Yong et al. (2010), even if as discussed above the size found for this complex can vary. In our case, the bands at ∼ 100 kDa and ∼ 150 kDa in Western-blots could thus correspond to incomplete denaturation of the protein complex at ∼ 480 kDa detected when doing BN-PAGE and/or residual disulfide bonding between two or three NRT2.1 monomers and/or other proteins different from NAR2.1. Indeed, in all the experiments we performed, NAR2.1 antibody failed to detect any band at ∼ 100 kDa and ∼ 150 kDa in Western blot (data not shown). Interestingly, the detection of these higher molecular weight bands revealed that the activation of NRT2.1 after 4h of NO_3_^-^ re-supply seems to be associated with the removal of NRT2.1 S501 phosphorylated form from the complex at ∼ 150 kDa and its appearance as a monomeric form at ∼ 45 kDa (Figure 7A). It explains why no apparent changes in the level of the overall NRT2.1 501 phosphorylated form was detected in ELISA (Figure 7B) and it reinforces our hypothesis that NRT2.1 activation is due to the synthesis of new unphosphorylated NRT2.1 protein and not to S501 dephosphorylation. Furthermore, it suggests that NRT2.1 activity depends on the composition of the high molecular weight complex, which would mainly contain NRT2.1 S501 phosphorylated form when it is inactive and the newly synthetized unphosphorylated form of NRT2.1 when it becomes active.

Considered together these results are consistent with a model (Figure 9) in which, (i) under no N conditions, NRT2.1 is phosphorylated on S501 residue and lead to the inactivation of a protein complex composed of several subunits of each NAR2.1 and NRT2.1 phosphorylated form, maybe associated with other unknown proteins and (ii) after addition of NO_3_^-^, NRT2.1 synthesis is induced and S501 phosphorylation is inhibited, leading to the activation of a protein complex composed of several subunits of each NAR2.1 and newly synthetized NRT2.1 unphosphorylated form, maybe associated with other unknown proteins.

**Figure 9.**
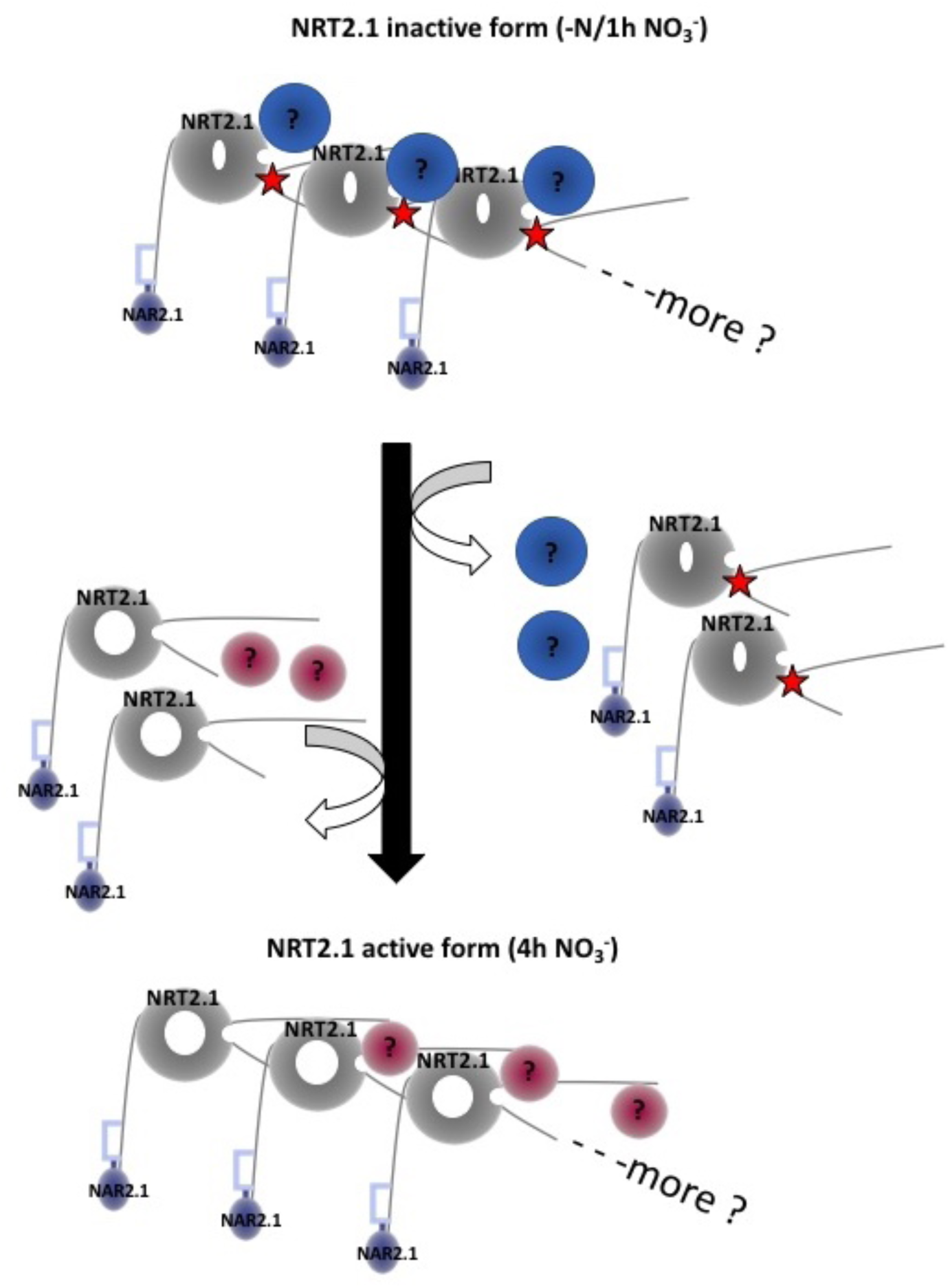
Model for the regulation of NRT2.1 activity NRT2.1 is represented with NAR2.1 interacting with its N-terminus and the red stars indicate S501 phosphorylation in NRT2.1 C-terminus part. The blue and pink disks represent hypothetical partner proteins, which could respectively participate in the inactivation and the activation of NRT2.1 protein complex. Under non inductive conditions, S501 is phosphorylated, the pores of NRT2.1 are closed and the protein complex is inactive. On inductive conditions, the protein complex with the phosphorylated form of NRT2.1 is dissociated and replaced by a protein complex containing new forms of non-phosphorylated NRT2.1. The absence of NRT2.1 S501 phosphorylation in the protein complex open the pores and enable NO_3_^-^ to enter the root.

### Impact of NRT2.1 S501 phosphorylation on root development

Beyond the role of NRT2.1 in root NO_3_^-^ uptake there are some indications that NRT2.1 could also act as a NO_3_^-^ sensor to coordinate the development of the root system with NO_3_^-^ availability. The first indication comes from a genetic screen, which isolated the *lin1* mutant resistant to the repressive effect of a high sucrose/NO_3_^-^ ratio on LR initiation (Malamy and Ryan, 2001). The *lin1* mutant was found to carry a missense mutation in *NRT2.1* and indicates that NRT2.1 acts as a repressor of LR initiation under high sucrose/low NO_3_^-^ supply (Little et al., 2005). The second indication comes from Remans et al. (2006), who also found a LR initiation phenotype for a *nrt2.1* mutant. However, in this study, *nrt2.1* plants initiated less LR primordia than the wild type under NO_3_^-^ limited condition suggesting an activator role for NRT2.1 in LR initiation. Despite these discrepancies, LR initiation phenotype of the *nrt2.1* mutants is observed in both studies even in the absence of added NO_3_^-^ in the external medium. Altogether, these data show that NRT2.1 seems to fulfill a dual transport/signaling role, thus displaying functional properties of a putative transceptor like for example the ammonium transporter MEP2 in yeast or the NO_3_^-^ transporter NRT1.1 in Arabidopsis (Lorenz and Heitman, 1998; Ho et al., 2009; Krouk et al., 2010).

One of the most convincing evidence for a membrane transporter to act as a transceptor is the genetic uncoupling of transport function and signaling effects. For example, under conditions of nitrogen limitation, MEP2 initiates a signaling cascade that triggers filamentous (pseudohyphal) growth of the yeast colonies. This response can be uncoupled from the ammonium transport function of MEP2 by point mutations within the central loop of the protein (Van Nuland et al., 2006). For NRT1.1, a point mutation, P492L, uncouples transport and sensing and in the corresponding mutant plant (*chl1-9*), transport is impaired, while the sensing function, namely the induction of *NRT2.1* expression, is still functional (Ho et al., 2009). For NRT2.1, our data show that S501D substitution does not allow uncoupling the transport function from the signaling effect. Indeed, S501D plants are impaired for root NO_3_^-^ uptake and have the same defect in LR initiation than *nrt2.1* mutant under low NO_3_^-^ conditions, while S501A plants behave like the WT for both NO_3_^-^ uptake and root development (Figure 8). It indicates that S501 phosphorylation does not directs the action of NRT2.1 towards the activation of specific responses. Since the defect in LR initiation in S501D plants is also observed on a N-free medium, it confirms, as described previously, that the signaling mechanism cannot be explained by a lack of NRT2.1 capacity to transport NO_3_^-^ (Little et al., 2005; Remans et al., 2006). Furthermore, it suggests that NRT2.1 signaling mechanism does not depend on the presence or absence of the protein since the same decrease in LR initiation is observed in both *nrt2.1* mutant and S501D plants (Figure 8). However, the mechanisms involved remain unknown and will require further investigation to elucidate the role of S501 phosphorylation in NRT2.1 NO_3_^-^ uptake activity and signaling for LR development.

### Conservation of NRT2.1 S501 phosphorylation site across plant species

A major role of S501 phosphorylation site for NRT2.1 activity is also supported by the fact that this residue is very strongly conserved in all Arabidopsis ecotypes (data not shown) and in all plant species where a clear NRT2.1 homolog has been identified (Figure 10). The alignment of NRT2.1 C-terminus reveals that except in yeast and fungus, the serine corresponding to S501 in Arabidopsis is remarkably conserved across most algae, mosses, dicotyledons and monocotyledons. The fact that S501 does not align with NRT2.1 homologs from fungus and yeast corresponds to previous models for the membrane topology of these polypeptides showing that NRT2.1 C-terminal extension is a general feature of NRT2 transporters from algae and higher plants, which is absent in yeast and fungus (Forde, 2000; Jacquot et al., 2017). However, more surprisingly, the serine corresponding to S501 in Arabidopsis is also noticeably replaced by a glycine in the three monocotyledons wheat (*Triticum aestivum*), barley (*Hordeum vulgare*) and *Brachypodium distachyon*, while it is conserved in rice for example (*Oryza sativa*). If there is for now no elements to explain this discrepancy between these three species and the other plants, it is interesting to note that such specificity for barley has already been observed for another residue, Ser463. Indeed, this residue also located in the C-terminus of HvNRT2.1, has been shown to be required for the interaction between HvNRT2.1 and HvNAR2.1 (Ishikawa et al., 2009). However, this residue is only conserved in NRT2.1 transporters from algae and monocotyledons, but not in dicotyledons (Jacquot et al., 2017). These discrepancies between essential amino acids for either NRT2.1 activity and/or interaction with partner proteins among different plant species, could reveal the existence of several molecular mechanisms for the regulation of NRT2.1 at the post-translational level. It seems to be at least already the case for the interaction between NAR2.1/NRT2.1, which appears to involve the C-terminus part of NRT2.1 for barley (Ishikawa et al., 2009) instead of the N-terminus part of NRT2.1 in Arabidopsis (Kotur et al., 2017).

**Figure 10.**
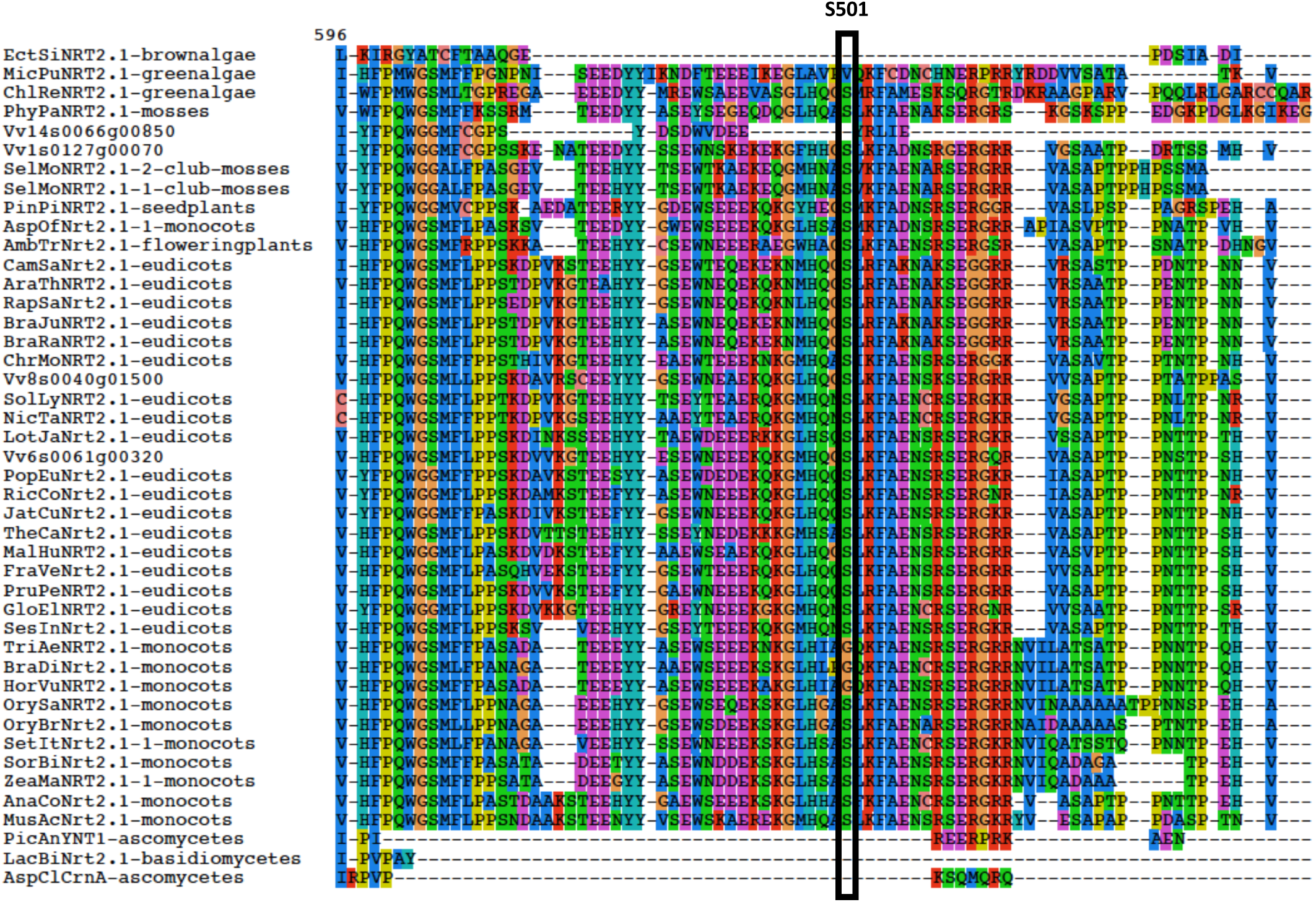
NRT2.1 C-terminus part alignment performed using Muscle v3.8.31 (Edgar, 2004) and displayed with SeaView version 4 (Gouy et al., 2010).

## Methods

### Generation of transformant lines

All constructs were made using Gateway technology (Invitrogen) according to the manufacturer’s instructions. For the truncated forms of NRT2.1, the *NRT2.1* DNA sequence from the *bac* clone T6D22 (ARBC, Columbus, OH), including the 1335-bp 5’ untranslated region and promoting sequence upstream the ATG, was amplified using the primers: NRT2.1 GATE forward 5’-CACCCACGTCAGCGAGATTGATCG-3’; NRT2.1 SANS reverse: 5’ TCACTGCTTCTCCTGCTCATTCCACTC 3’ (ΔC_494-530_ plants) or NRT2.1 AVEC reverse: 5’ TCAGCGTCCACCCTCTGACTTGGCGTT 3’ (ΔC_514-530_ plants). For site directed mutagenesis the *NRT2.1* DNA sequence from the *bac* clone T6D22 was amplified using: NRT2.1 GATE forward 5’-CACCCACGTCAGCGAGATTGATCG-3’; GATE NRT2.1 R, 5’-TCAAACATTGTTGGGTGTGTT-3’. Each amplification was cloned into the pENTR™/D-Topo vector (Invitrogen) to create entry clones. For site directed mutagenesis, NRT2.1 mutations were introduced by PCR of the whole entry clone with primers pair containing the desired site-directed mutation (NRT2.1 S501A Fw 5’-GAACATGCATCAAGGAGCTCTCCGGTTTGCCGAG-3’, NRT2.1 S501A Rev 5’-CTCGGCAAACCGGAGAGCTCCTTGATGCATGTTC-3’; NRT2.1 S501D Fw 5’-GAACATGCATCAAGGAGATCTCCGGTTTGCCGAG-3’, NRT2.1 S501D Rev 5’-CTCGGCAAACCGGAGATCTCCTTGATGCATGTTC-3’). After transformation of MacCell™ DH5*α* 10^9^ thermocompetent *Escherichia coli* (iNtRON Biotechnology), clones were fully sequenced before subsequent cloning in the binary Gateway destination vector pGWB501 obtained from Tsuyoshi Nakagawa (Research Institute of Molecular Genetics, Shimane University, Matsue, Japan) by using a Gateway LR Clonase enzyme mix (Invitrogen). The binary construct was introduced into the *Agrobacterium tumefaciens* strain GV3101, and the resulting bacterial culture was used to transform the *atnrt2.1-2* mutant plants, ecotype Col-0, by the standard flower dip method (Clough and Bent, 1998). The transformants were selected on a medium containing hygromycin (25 µg.mL^-1^). For further analyses, T1 segregation ratios were analyzed to select transformants with one T-DNA insertion and to isolate T3-homozygous plants.

### Growth conditions

Plants were grown hydroponically using the experimental set-up described previously (Lejay et al., 1999). Briefly, seeds were sown directly on the surface of wet sand in modified 1.5-ml microcentrifuge tubes, with the bottom replaced by a metal screen. The tubes supporting the seeds were placed on polystyrene floating rafts, on the surface of a 10-liter tank filled with tap water. The culture was then performed in a controlled growth chamber with 8 H/16 H day/night cycle at 23/20 °C and RH of 65 %. Light intensity during the light period was at 250 µmol.m^-2^.s^-1^. One week after sowing, the tap water was replaced by nutrient solution until the age of 5– 7 weeks depending on the size of the plants. The basal nutrient solutions supplied to the plants are those described by (Gansel et al., 2001) and contained either 1 mM NO_3_^-^ or 10 mM NH_4_NO_3_as nitrogen sources. The nutrient solution was replaced every week during this period and the day before the experiment. For NO_3_^-^ induction experiments, plants grown on 1 mM NO_3_^-^ were transferred during 5 days on a N-free solution before the experiment.

For root architecture analysis, the plants were grown in vertical agar plates on a 1 mM NO_3_^-^ in vitro culture as described in Remans et al. (2006). The culture was performed in a controlled growth chamber with 16 H/8 H day/night cycle at 20/20 °C and RH of 70 %. Light intensity during the light period was at 250 µmol.m^-2^.s^-1^. After 6 days of development, the seedlings were transferred to 0 mM NO_3_^-^, 0.3 mM NO_3_^-^ (LN) or 5 mM NO_3_^-^ (HN). The plants were then harvested after 7 days of development. Length measurements of the main root were performed on the Fiji® software. Quantification of primordia was performed with an OLYMPUS BH2 microscope. The average density of primordia was obtained in relation to the size of the primary root.

### RNA Extraction and Gene Expression Analysis

Root samples were frozen in liquid N_2_ in 2-mL tubes containing one steel bead (2.5 mm diameter). Tissues were disrupted for 1 min at 28 s^-1^ in a Retsch mixer mill MM301 homogenizer (Retsch, Haan, Germany). Total RNA was extracted from tissues using TRIzol reagent (Invitrogen, Carlsbad, CA, USA). Subsequently, 4 µg of RNAs were treated with DNase (DNase I Amplification Grade, Sigma) following the manufacturer’s instructions. Reverse transcription was achieved with 4 µg of RNAs in the presence of Moloney Murine Leukemia Virus (M-MLV) reverse transcriptase, (RNase H minus, Point Mutant, Promega) after annealing with an anchored oligo(dT)_18_ primer as described by Wirth et al. (2007). The quality of the cDNA was verified by PCR using specific primers spanning an intron in the gene *APTR* (At1g27450) forward 5’-CGCTTCTTCTCGACACTGAG-3’; reverse 5’-CAGGTAGCTTCTTGGGCTTC-3’.

Gene expression was determined by quantitative real-time PCR (LightCycler 480, Roche Diagnostics) using the SYBR^R^ Premix Ex Taq™ (TaKaRa) according to the manufacturer’s instructions with 1 µl of cDNA in a total volume of 10 µl. The conditions of amplifications were performed as described previously by Wirth et al. (2007), except the first 10 minutes at 95°C which has become 30 seconds at 95°C. All the results presented were standardized using the housekeeping gene Clathrin (At4g24550). Gene-specific primer sequences were: NRT2.1 forward, 5’-AACAAGGGCTAACGTGGATG-3’; NRT2.1 reverse, 5’-CTGCTTCTCCTGCTCATTCC-3’; NAR2.1 forward, 5’-GGCCATGAAGTTGCCTATG-3’; NAR2.1 reverse, 5’-TCTTGGCCTTCCTCTTCTCA-3’; Clathrin forward, 5’-AGCATACACTGCGTGCAAAG-3’; Clathrin reverse, 5’-TCGCCTGTGTCACATATCTC-3’.

### NO_3_^-^ influx studies

Root NO_3_^-^ influx was assayed as described by (Delhon et al., 1995). Briefly, the plants were sequentially transferred to 0.1 mM CaSO_4_ for 1 min, to a complete nutrient solution, pH 5.8, containing 0.2 mM ^15^NO_3_ (99 atom % excess^15^N) for 5 min, and finally to 0.1 mM CaSO_4_ for 1 min. Roots were then separated from shoots, and the organs dried at 70 °C for 48 h. After determination of their dry weight, the samples were analyzed for total nitrogen and atom % ^15^N using a continuous flow isotope ratio mass spectrometer coupled with a C/N elemental analyzer (model Euroflash Eurovector, Pavia Italy) as described in (Clarkson, 1986).

### NRT2.1 Immunodetection and Membrane purification

Microsomes were prepared as described by (Giannini et al., 1987) and plasma membrane vesicles were purified from microsomes by aqueous two-phase partitioning, as described by (Santoni et al., 2003).

Proteins were separated on denaturing SDS-PAGE followed by an electrotransfer at 4 °C onto a PVDF membrane (0.2 µM, Bio-Rad). NRT2.1 was detected using three different anti-NRT2.1 antisera produced by Eurogentec against either the synthetic peptide TLEKAGEVAKDKFGK (anti-NRT2.1 19), CKNMHQGSLRFAENAK (anti-NRT2.1 20) or KNMHQG(p)SLRFAENAK (anti-S501P). NAR2.1 was detected using one anti-NAR2.1 antisera produced by Eurogentec against the synthetic peptide DVTTKPSREGPGVVL (anti-NAR2.1). All the polyclonal antisera were affinity purified by Eurogentec. The immunodetection was performed with a chemiluminescent detection system kit (Pierce™ ECL Western Blotting Substrate, Pierce).

For ELISA, serial 2-fold dilutions in a carbonate buffer (30 mM Na_2_CO_3_, 60 mM NaHCO_3_, pH 9.5) of 20 µg of microsomes proteins were loaded in duplicate on Maxisorb immunoplates (Nunc, Denmark) and left overnight at 4°C. The immunodetection was performed according to the manufacturer’s instructions. The primary anti-NRT2.1 20 (1:2500 dilution) and anti-S501P antibodies (1:500 dilution) and a secondary peroxidase-coupled anti-rabbit antibody were successively applied for 2 h at 37°C. A linear regression between the absorbance signal due to oxidized 2,2’-azinobis-3-ethylbenzothiazolinz-6-sulfonic acid diammonium salt, as read with a multiplate reader (Victor, PerkinElmer Life sciences), ant the amount of proteins was obtained for each sample and used for relative comparison between samples.

For Dot blot analysis, the synthetic peptides, CKNMHQGSLRFAE and CKNMHQG(p)SLRFAE, produced by Eurogentec for the purification of the antibody anti-S501P, were spotted onto a PVDF membrane (0.45 µM, Hybond-P, Amersham). The membrane was then left to dry for 1.5 h at room temperature before being probed with S501-P antibody or the purified antibody produced against the non-modified peptide KNMHQGSLRFAENAK.

Blue Native PAGE (BN-PAGE) was adapted from (Peltier et al., 2001; Peltier et al., 2004) and (Kotur and Glass, 2015). A volume of plasma membrane (PM) resuspension buffer containing 10% dodecyl *β*-D-maltoside (DDM) was added to PM suspension to have 1.5% DDM in the final solution. After disruption of the samples with a potter and incubation at 4°C for 15 min, solubilized samples were combined with 1/10 volume of sample loading buffer (5% Serva Blue G in 50 mM BisTris/HCl (pH 7.2), 5 mM MgCl_2_, 0.75 M 6-amino-N-caproic acid, 20% w/v glycerol, 10% v/v protease inhibitors cocktail). After a second incubation at 4°C for 15 min, samples were applied to 1.5 mm thick 16×16 cm native gradient gels (5–16% acrylamide) in vertical electrophoresis unit operated at 4°C. Gel lanes were then cut and the proteins were denaturated, reduced, and alkylated in SDS loading buffer (50 mM Tris pH 6.8, Urea 6 M, glycerol 30% and SDS 2%) containing either 100 mM DTT for the first 30 min and 260 mM iodoacetamid for another 30 min. For separation in the second dimension, the gel lanes were then placed into a 12% acrylamide Tricine 16×24 cm gel of the same thickness in vertical electrophoresis unit operated at room temperature and then transferred on a PVDF membrane (0.2 µM, Bio-Rad) for immunodetection.

### Mass Spectrometry

Microsomes of both wild-type Col-0 ecotype and GFP10 transgenic plants (Wirth et al., 2007) grown in hydroponics were used to determine NRT2.1 phosphorylation sites.

For S11, S28 and S501, the filter-aided sample preparation protocol (Wisniewski et al., 2009) was used to perform in solution reductions/alkylations simultaneously with detergent removing. The proteins were then subsequently digested at 37°C with Lys-C (Roche Applied Science) during 4 h then with trypsin (Sequencing Grade Modified, Promega, Madison, WI) overnight. Peptides were eluted by step elutions with 50 mM ammonium bicarbonate, followed by a 50% ACN and then 0.5 M NaCl elution steps. Peptides were desalted on C18 columns (Sep-Pak VactC18 cartridge 1cc, Waters). For Mass Spectrometry peptides were then concentrated with a precolumn (ThermoScientific, C18 PepMap100, 300 µm × 5 mm, 5 µm, 100 Å) and separated with a reversed-phase capillary column (ThermoScientific, C18 PepMap100, 75 µm × 250 mm, 3 µm, 100 Å) over a 90 min gradient (300nL/min). Peptide fragmentation was carried out with a ion trap tandem MS system (AmaZon Speed ETD, Bruker) with nanoSprayer (Bruker), performed in the positive ion mode and alternating collision induced dissociation (CID) and electron transfer dissociation (ETD). MS and MS/MS spectra were acquired with a mass range of m/z 300-1500 in enhanced resolution (FWHM 0,3u). ICC target was fixed to 4×10^4^ ions with a maximum ion accumulation time set to 200 ms. The corresponding data was processed via DataAnalysis 4.4 (Bruker). Data was interrogated via ProteinScape 4.0 (Bruker) against SwissProt restricted to *Arabidopsis thaliana* (thale cress) taxonomy using an in-house Mascot search server (Matrix Science; version 2.6). The selected enzyme was trypsin. Parameters of interrogation accepted one putative missed cleavage, a 0,8 Da and 2 Da mass range for the parent peptide and for MS/MS fragment, respectively. Since proteins were reduced and alkylated, carbamidomethylation was selected as a fixed modification. Variable modifications were phosphorylation of serines, threonines, and tyrosines; oxidation of methionine; and acetylation of N-term proteins. For each analysis, peptides were determined by significant threshold p < 0.05.

For T521, immuno-purification was performed with µMACS GFP Tag Protein Isolation kits (Miltenyi Biotec). 1 mg of solubilized microsomes were loaded on µcolumns. Kit protocol was followed. Proteins were eluted with elution buffer contained 50 mM Tris HCl (pH 6.8), 50 mM DTT, 1% SDS, 1 mM EDTA, 0.005% bromophenol blue, 10% glycerol and conserved at - 20°C. The samples were then treated as described above except for peptide fragmentation which was carried out with a orbitrap MS system (LTQ-Orbitrap XL-ETD, ThermoScientific) with nanoSource (ThermoScientific), performed in the positive ion mode and alternating CID / ETD fragmentation. The corresponding data was processed via Xcalibur 2.0.7 (ThermoScientific) with the 3 most intense precursors, bi-charged with alternating CID/ETD fragmentation. Data was interrogated against a local NRT2.1-GFP database using an in-house Mascot search server (Matrix Science; version 2.4) with the same parameters as for S11, S28 and S501.

### rBiFC

Open reading frames of *NRT2.1* and *NAR2.1* were amplified with gene-specific primers that included Gateway attachment sites (attB3/B2 or attB1/B4). Subsequent BP reactions in pDONR221-P3P2, and pDONR221-P1P4 (Invitrogen) yielded Entry clones that were verified via sequencing. *NRT2.1* and *NAR2.1* sequences were obtained without stop codon to allow C-terminal fusions. Gateway Destination clones were generated using LR Clonase II (Invitrogen) by LR reaction according to the manufacturer’s instructions.

Point mutants were generated by site-directed mutagenesis kit (Agilent Technologies). Primers of different point mutants were designed by online tool QuikChange Primer Design (Agilent Technologies). All primers used for this experiment are listed in Table 1.

**Table 1:**
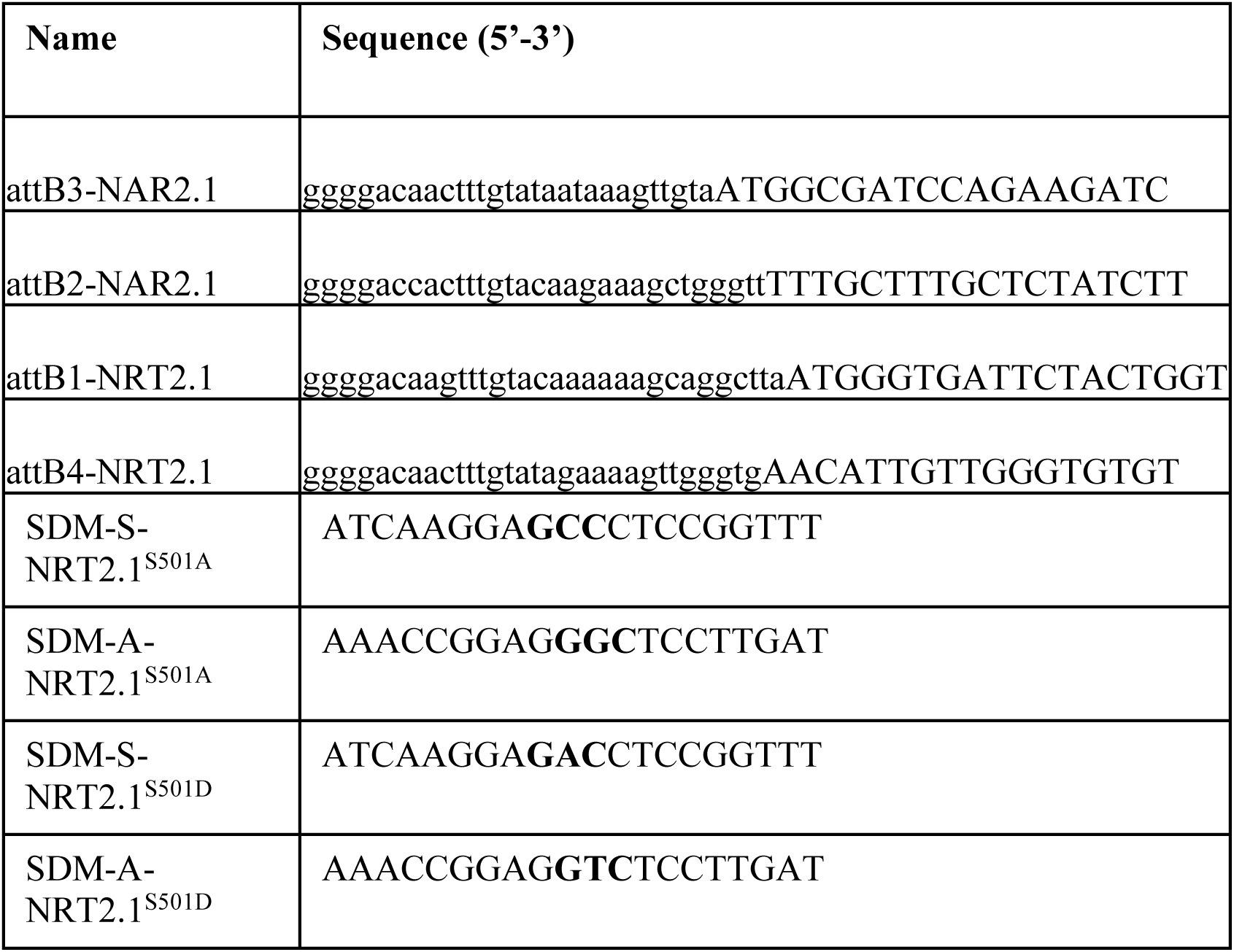
Oligonucleotides used for Entry clone design and site-directed mutagenesis (SDM)

*Agrobacterium tumefaciens* strain GV3101 was used to transiently transform *Nicotiana benthamiana* with constructs as described previously through syringe mediated infiltration (Schob et al., 1997; Sparkes et al., 2006). Overnight cultures of agrobacteria were used to inoculate fresh LB medium (supplemented with 50 µg/mL rifampicin, and 100 µg/mL spectinomycin) and grown for another 4 to 6 h up to an optical density at 600 nm of 0.6 to 0.7. Agrobacteria were harvested, and resuspended in AS medium (10 mM MgCl2, 10 mM MES-KOH, pH 5.6, plus 150 mM acetosyringone (Grefen et al., 2008; Blatt and Grefen, 2014)). After an incubation time of 60 min at room temperature, leaves of 4-week-old *Nicotiana benthamiana* were infiltrated as described previously (Schob et al., 1997; Sparkes et al., 2006; Blatt and Grefen, 2014). Leaves were subjected to CLSM analysis 2 d after infiltration.

For rBiFC assays, confocal images were collected using a Zeiss LSM700 confocal microscope with 20X/0.75-NA objectives. Excitation intensities, filter settings, and photomultiplier gains were standardized. YFP at excitation 514-nm and emission 521- to 565-nm, RFP at excitation 545-nm and emission 560- to 615-nm.

rBiFC fluorescence ratios were calculated as a ratio of YFP and RFP fluorescence described previously (Grefen and Blatt, 2012).

## Acknowledgements

The work was supported by an international grant from the ANR in France and DFG in Germany (SIPHON ANR-13-ISV6-0002-01), by a national grant from the ANR (TransN ANR-BLAN-NT09_477214) and by post-doctoral funding from the CNRS (E.L.).

## Author contributions

AJ performed most of the experiments including the generation and the characterisation of the transgenic plants, Western Blots and BN-PAGE. VC and LL obtained and performed experiments with the antibody anti-S501P and participated to BN-PAGE experiments. VC, FB and AnM performed root architecture experiments. AdM performed mass spectrometry experiments to find NRT2.1 phosphorylation sites along with VS and SH. AdM and EL optimised BN-PAGE protocol for NRT2.1. ZL and WS performed rBiFC experiments. PT performed ^15^N measurements, CF performed NRT2.1 sequence analysis and PB participated to the characterisation of the transgenic plants. LL and WS conceived the project and designed the experiments. LL, AG and WS interpreted the data and wrote the manuscript.

---

**Supplemental Figure 1.** Root NO3- influx in response to sucrose, light and NH4N03 in the transgenic lines ΔC_494-530_, ΔC_514-530_, S501A and S501D.

(A) Plants were grown on 1 mM NO_3_^-^ and after a normal night were either kept in the dark during 4h on a complete nutrient solution with or without 1% sucrose or transferred during 4h in the light.

(B) Plants were grown on 1 mM NO_3_^-^ and were transferred during 4h on a solution containing 10 mM NH_4_NO_3_.

Root NO_3_^-^ influx was measured at the external concentration of 0.2 mM ^15^NO_3_^-^. Values are means of 12 replicates ± SD.

**Supplemental Figure 2.** rBiFC analysis for NRT2.1 and NAR2.1 interaction.

rBiFC analysis of YFP and RFP fluorescence collected from tobacco plants transformed using the pBiFCt-2in1-CC vector. Left to right, images are YFP (BiFC) fluorescence, RFP fluoresence and bright field. Bar=50µm

**Supplemental Figure 3.** Dot blot analysis for the specificity of the antibody Anti-S501P.

Serial 2 fold dilution of 3 µg of synthetic peptides phosphorylated or not on S501 were blotted. Membranes were probed with either the antibody specific for S501 phosphorylation (Anti-S501P) or the antibody non-specific to the modification.

**Supplemental Figure 4.** Root NO3- influx after 1h and 4h of NO_3_^-^ induction.

Wild type (Col) and *nrt2.1-2* knockout mutant (*nrt2.1*) were grown on 1 mM NO_3_^-^ for 5 weeks and were starved for N during 5 days. Thereafter, the plants were re-supplied with 1 mM NO_3_^-^ during 1h or 4h. Root NO_3_^-^ influx was measured at the external concentration of 0.2 mM ^15^NO_3_. Values are means of 12 replicates ± SD.

