## Supplemental Figure 1 for "NRT2.1 phosphorylation prevents root high affinity nitrate uptake activity in *Arabidopsis thaliana*"

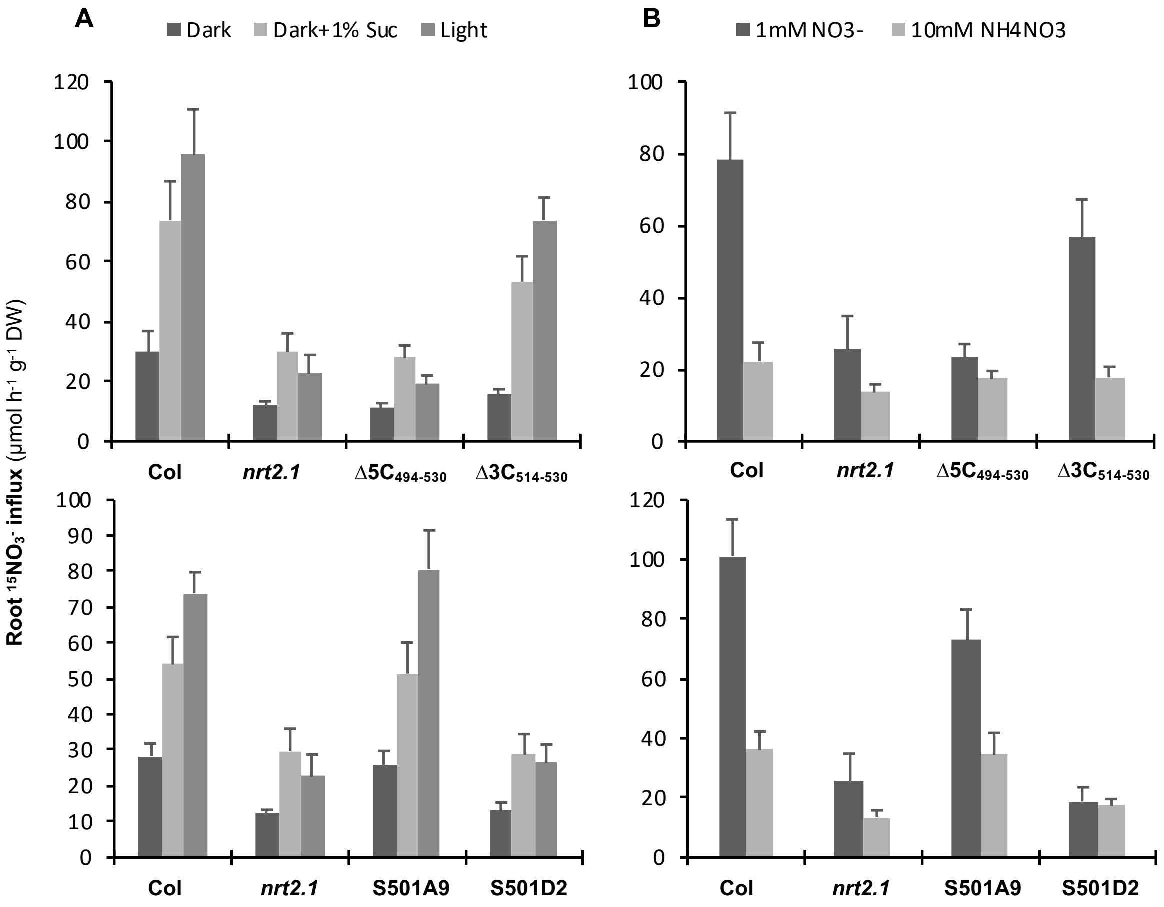


**Supplemental Figure 1.** Root $NO_3^-$ influx in response to sucrose, light and $NH_4N0_3$ in the transgenic lines $\Delta C_{494\text{-}530}$, $\Delta C_{514\text{-}530}$, S501A and S501D.

**(A)** Plants were grown on 1 mM $NO_3^-$ and after a normal night were either kept in the dark during 4h on a complete nutrient solution with or without 1% sucrose or transferred during 4h in the light.
**(B)** Plants were grown on 1 mM $NO_3^-$ and were transferred during 4h on a solution containing 10 mM $NH_4NO_3$.
Root $NO_3^-$ influx was measured at the external concentration of 0.2 mM $^{15}NO_3^-$. Values are means of 12 replicates ± SD.
