## Supplemental Figure 2 for "NRT2.1 phosphorylation prevents root high affinity nitrate uptake activity in *Arabidopsis thaliana*"

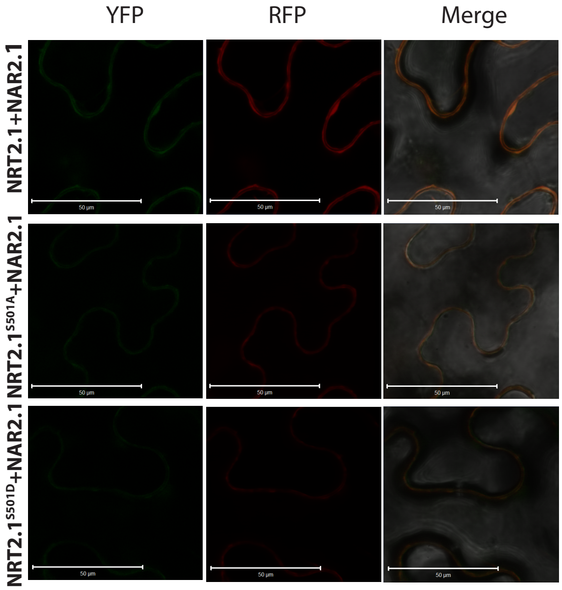


**Supplemental Figure 2.** rBiFC analysis for NRT2.1 and NAR2.1 interaction.

rBiFC analysis of YFP and RFP fluorescence collected from tobacco plants transformed using the pBiFCt-2in1-CC vector. Left to right, images are YFP (BiFC) fluorescence, RFP fluoresence and bright field. Bar=50µm
