## Supplemental Figure 3 for "NRT2.1 phosphorylation prevents root high affinity nitrate uptake activity in *Arabidopsis thaliana*"

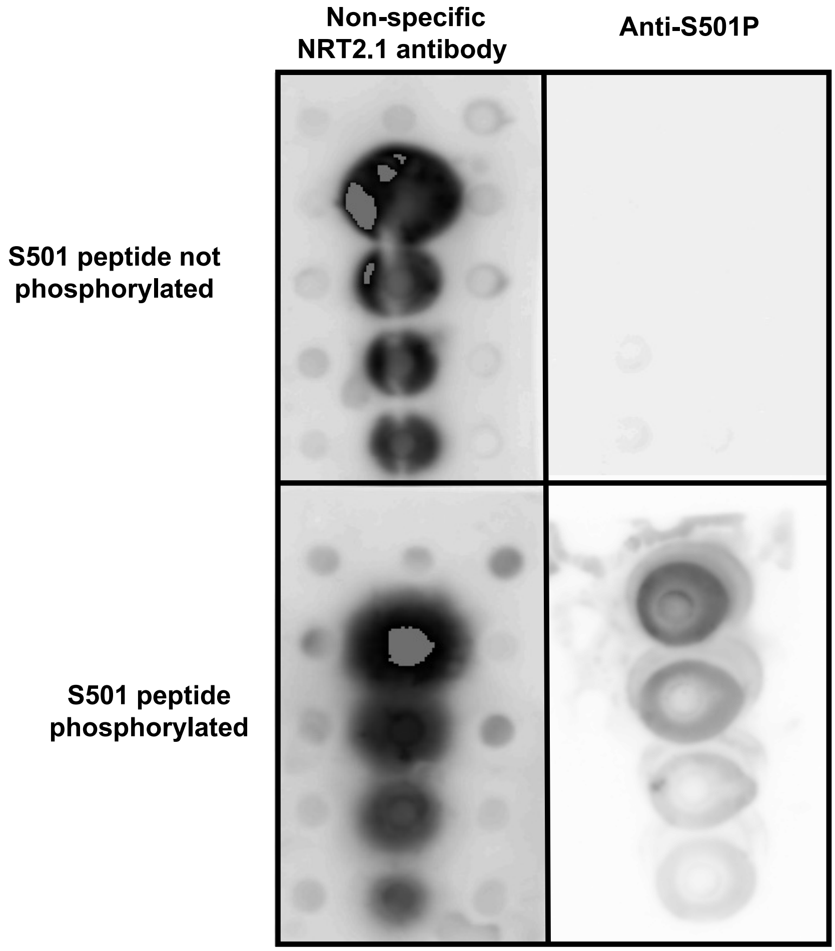


**Supplemental Figure 3.** Dot blot analysis for the specificity of the antibody Anti-S501P.

Serial 2 fold dilution of 3 µg of synthetic peptides phosphorylated or not on S501 were blotted. Membranes were probed with either the antibody specific for S501 phosphorylation (Anti-S501P) or the antibody non-specific to the modification.
