## Supplemental Figure 4 for "NRT2.1 phosphorylation prevents root high affinity nitrate uptake activity in *Arabidopsis thaliana*"

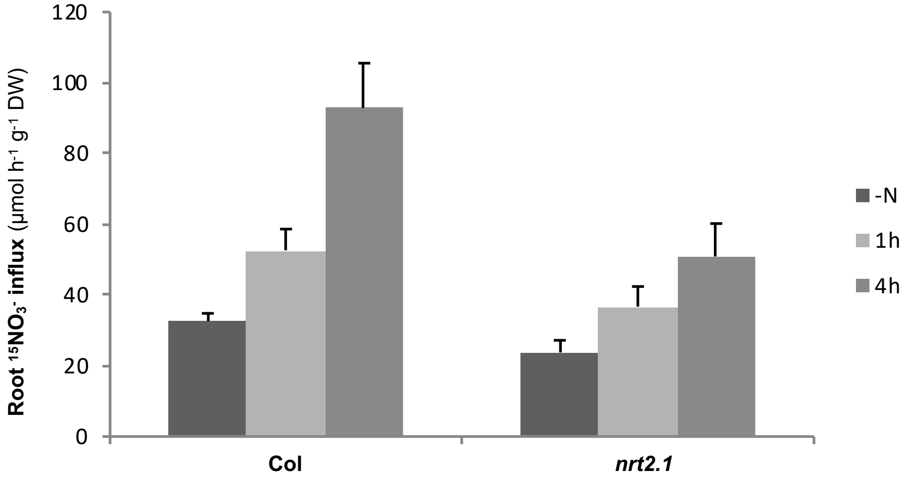


**Supplemental Figure 4.** Root $NO_3^-$ influx after 1h and 4h of $NO_3^-$ induction.

Wild type (Col) and *nrt2.1-2* knockout mutant (*nrt2.1*) were grown on 1 mM $NO_3^-$ for 5 weeks and were starved for N during 5 days. Thereafter, the plants were re-supplied with 1 mM $NO_3^-$ during 1h or 4h. Root $NO_3^-$ influx was measured at the external concentration of 0.2 mM $^{15}NO_3$. Values are means of 12 replicates ± SD.
